# SMAD4 and KRAS status shape malignant-stromal crosstalk in pancreatic cancer

**DOI:** 10.1101/2024.04.28.591518

**Authors:** Eloise G. Lloyd, Muntadher Jihad, Judhell S. Manansala, Wenlong Li, Priscilla S. W. Cheng, Sara Pinto Teles, Gianluca Mucciolo, Joaquín Araos Henríquez, Sally Ashworth, Weike Luo, Sneha Harish, Paul M. Johnson, Lisa Veghini, Marta Zaccaria, Rebecca Brais, Mireia Vallespinos, Vincenzo Corbo, Giulia Biffi

## Abstract

Pancreatic ductal adenocarcinoma (PDAC) contains an extensive stroma that modulates response to therapy, contributing to the dismal prognosis associated with this cancer. Evidence suggests that the stromal composition of PDAC is shaped by mutations within malignant cells; however, most pre-clinical models of PDAC are driven by *Kras^G12D^* and mutant *Trp53* and have not assessed the contribution of other known oncogenic drivers, including *KRAS^G12V^* and alterations in *CDKN2A* and *SMAD4*. To increase understanding of malignant cell-stroma crosstalk in PDAC, we analyzed *Trp53-*mutant mouse models driven by *Kras^G12D^* or *Kras^G12V^* in which *Smad4* was wild-type or deleted. *Kras^G12D^*; *Smad4*-deleted PDAC developed a fibro-inflammatory rich stroma with increased JAK/STAT malignant cell signaling and an enhanced therapeutic response to JAK/STAT inhibition. In stark contrast, the stroma of *Smad4*-deleted *Kras^G12V^* PDAC was differently altered, and the malignant compartment lacked JAK/STAT signaling dependency. Thus, malignant cell genotype impacts malignant-stromal phenotype in PDAC, directly affecting therapeutic efficacy.

**STATEMENT OF SIGNIFICANCE:** Understanding malignant cell-stroma crosstalk in PDAC has focused on models containing *Kras^G12D^* and mutant *Trp53*. Here, we show that PDAC driven by *Kras^G12D^* or *Kras^G12V^*in which *Smad4* is deleted display differences in malignant-stromal signaling and treatment sensitivity, highlighting the importance of understanding genotype-phenotype relationships for precision PDAC therapy.

## INTRODUCTION

Pancreatic ductal adenocarcinoma (PDAC) is the fourth most common cause of cancer-related death and >85% of patients succumb to their disease within 5 years (1). PDAC is characterized by an extensive stroma that contributes to this dismal prognosis. Cancer-associated fibroblasts (CAFs) are abundant stromal cells that have been shown to modulate PDAC progression and therapy response (2). Distinct populations of CAFs have been described in PDAC and other malignancies (2), including myofibroblastic CAFs (myCAFs), inflammatory CAFs (iCAFs) and antigen-presenting CAFs (apCAFs) (3–5). Additional CAF subsets have been identified within or across these CAF states with distinct cells of origin, malignant cell-mediated reprogramming and/or functions (2–4,6–12). In addition to CAFs, other stromal cell types, including macrophages and neutrophils, have been recently shown to be more heterogenous than previously appreciated (13,14). Thus, improving PDAC survival for all patients will likely require a better understanding of the heterogenous nature of the tumor microenvironment (TME) in these cancers (15). Determining malignant cell-stroma vulnerabilities in distinct patient-relevant contexts may guide the design of precision cancer therapies for PDAC.

PDAC can be classified as basal/squamous or classical/progenitor subtypes, which contain different TMEs (16–20). However, this classification has not yet translated into improved treatment design, perhaps because these two subtypes can co-exist (21–23). Identifying distinct groups of PDAC with specific malignant cell/stroma characteristics and therapeutic sensitivity thus remains a priority. Four main genetic drivers of PDAC have been described: *KRAS* mutations (present in >90% patients) - with G12D (>40% patients) and G12V (>30% patients) mutations being the most abundant, *TP53* mutations (>70% patients), *SMAD4* loss (>30% patients) and *CDKN2A* loss (>30% patients) (24,25). However, most current knowledge of stroma composition comes from KRAS^G12D^, p53 mutant KPC (*Kras^LSL-G12D/+^*; *Trp53 ^LSL-R172H/+^*; *Pdx1*-*Cre*) genetically engineered mouse models (GEMMs) (26), which largely recapitulate patient disease progression but may not capture stromal differences across human PDAC. Indeed, evidence indicates that mutations in PDAC malignant cells can differently shape the stroma (27–30). For example, increased fibrosis and reduced CD8^+^ T cell infiltration has been associated with *TP53*-mutated PDAC, while *BRCA*-mutated PDAC were found to have different CAF composition compared to *BRCA*-WT tumors (27,28). Moreover, the stroma affects PDAC progression and therapy response (2). Therefore, distinct groups of PDAC may respond differently to therapies depending on their genotype and associated TME.

Malignant cell TGF-β signaling is a key player in shaping the PDAC stroma since it drives myCAF formation (5,11). Loss of *SMAD4*, which is involved in TGF-β signaling, is typically a late event in PDAC development and correlates with metastasis formation and worse prognosis (25,31). This effect may be in part dependent on the ability of TGF-β signaling-deficient malignant cells to remodel the epithelial and stromal compartments (30). To determine how SMAD4 deficiency tunes the PDAC TME and identify candidate malignant cell-stroma dependencies in this group of aggressive tumors, we established *Trp53-*mutant mouse models driven by *Kras^G12D^* or *Kras^G12V^* in which *Smad4* was wild-type or deleted.

## RESULTS

### *Smad4* loss impacts the immune TME in KPC PDAC

Human PDAC tumors display remarkable inter-tumoral clinical, histological, and genetic heterogeneity, rendering it difficult to understand how mutations cooperate to drive tumorigenesis (**Supplementary Fig. S1A-B**; **Supplementary Table S1**). Therefore, to start to better dissect the relationship between malignant cell genotype and stromal phenotype, we engrafted patient-derived PDAC organoids harboring either the *KRAS^G12D^* (KP) or *KRAS^G12V^* (KvP) mutation and *SMAD4* WT or deficient status into the pancreata of NOD SCID gamma (NSG) mice (33) (**Fig. 1A**; **Supplementary Fig. S1C-D** and **Supplementary Table S2**). SMAD4 loss significantly decreased the desmoplasia – measured using collagen deposition and alpha smooth muscle actin (αSMA) levels – and accelerated PDAC progression in both KP and KvP *TP53-*mutant PDAC (**Fig. 1B-F**; **Supplementary Fig. S1E-G**).

**Figure 1.**
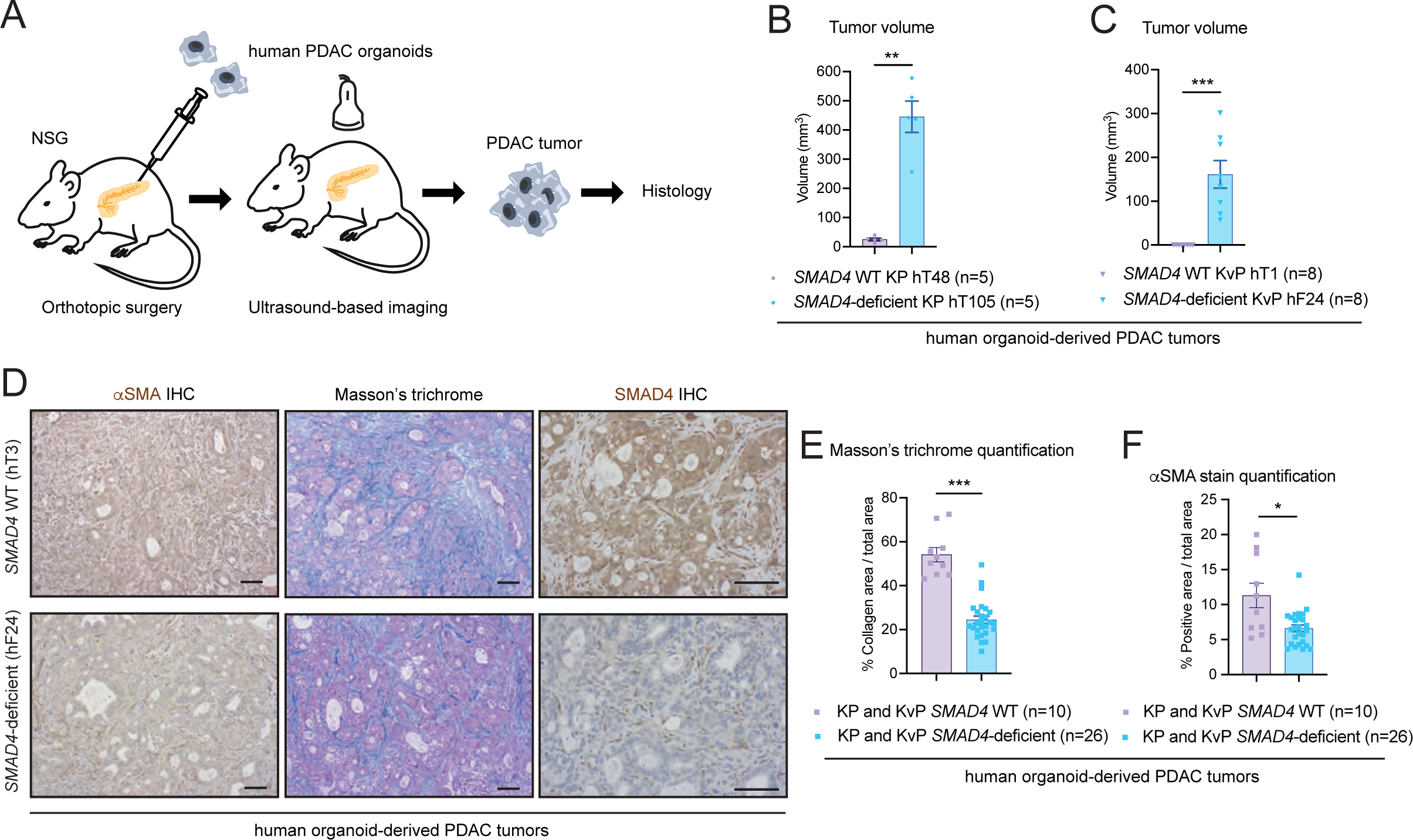
*SMAD4*-deficient human organoid-derived PDAC tumors have less fibrosis than *SMAD4* WT tumors. **(A)** Schematic of analyses of orthotopically-grafted human organoid-derived pancreatic ductal adenocarcinoma (PDAC) models in NOD SCID gamma (NSG) mice. **(B-C)** Tumor volumes as measured by ultrasound-based imaging of tumors derived from the transplantation of KP **(B)** or KvP **(C)** *SMAD4* wild-type (WT) or *SMAD4*-deficient human PDAC organoids with *KRAS*^G12D^ or *KRAS*^G12V^ mutation, respectively. Results show mean ± SEM. **, *P* < 0.01; ***, *P* < 0.001, Mann-Whitney test. **(D)** Representative SMAD4, Masson’s trichrome and alpha smooth muscle actin (αSMA) stains in *SMAD4* WT or *SMAD4*-deficient human organoid-derived PDAC tumors. Scale bars, 50 μm. **(E)** Quantification of Masson’s trichrome stain in *SMAD4* WT or *SMAD4*-deficient human organoid-derived PDAC tumors. Results show mean ± SEM. ***, *P* < 0.001, Mann-Whitney test. **(F)** Quantification of αSMA stain in *SMAD4* WT or *SMAD4*-deficient human organoid-derived PDAC tumors. Results show mean ± SEM. *, *P* < 0.05, Mann-Whitney test.

*KRAS^G12D^* is the most frequent *KRAS* mutant allele in PDAC patients (25). Thus, to further study the role of KRAS and SMAD4 status in PDAC, we deleted *Smad4* from KPC (*Kras^LSL-G12D/+^*; *Trp53 ^LSL-R172H/+^*; *Pdx1*-*Cre*) GEMM-derived organoids (26,34,35) (**Supplementary Fig. S2A-D**). Isogenic *Smad4* knockout (KO) KPC (hereafter, KPC*^Smad4-^*^KO^) organoids grew significantly faster that *Smad4* WT KPC (hereafter, KPC*^Smad4^*^-WT^) organoids both *in vitro* and as orthotopically grafted tumors in immunocompromised nu/nu or immunocompetent C57BL/6J mice (**Fig. 2A-C**; **Supplementary Fig. S2E-G**).

**Figure 2.**
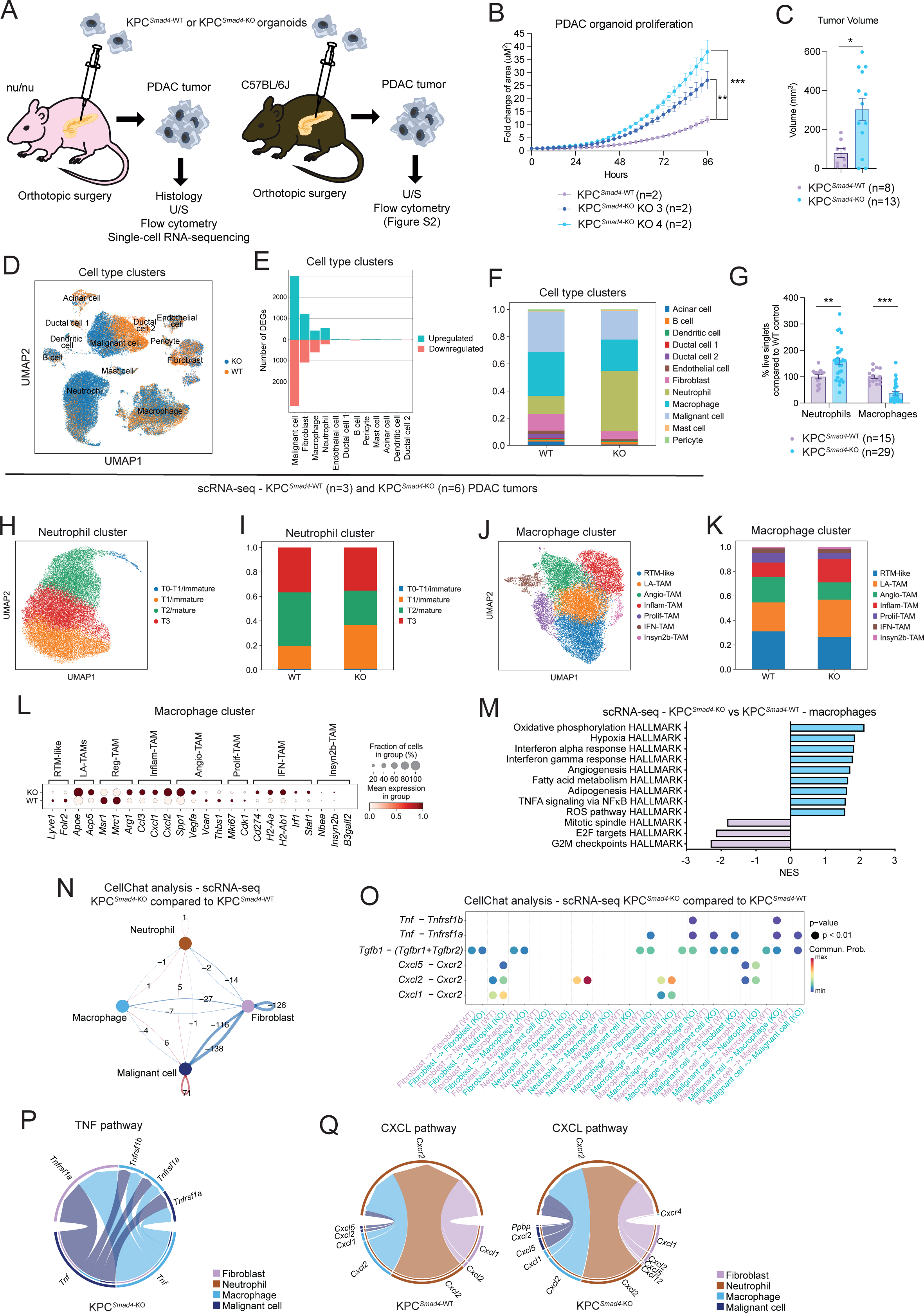
*Smad4* loss impacts the immune TME in KPC PDAC. **(A)** Schematic of analyses of KPC (i.e. *Kras*^G12D^ p53 mutant) organoid-derived PDAC models in nu/nu (left) or C57BL/6J (right) mice. **(B)** Proliferation curves of KPC*^Smad4^*^-WT^ or KPC*^Smad4^*^-KO^ PDAC organoids cultured for 96 hours in Matrigel in organoid complete media. Data were normalized to the first measurement (at 3 hours post-plating on day 0). Results show mean ± SEM of n=2 biological replicates (with n=4 technical replicates each). **, *P* < 0.01; ***, *P* < 0.001, Mann-Whitney test calculated for the last time point. **(C)** Tumor volumes as measured by ultrasound-based imaging of tumors derived from the transplantation of KPC*^Smad4^*^-WT^ or KPC*^Smad4^*^-KO^ PDAC organoids in nu/nu mice. Results show mean ± SEM from 2 separate experiments, each with 1 WT group and 2 groups of KO pools from 2 different guides (n=8-13 mice/cohort; at day 25 (experiment 1) or day 21 (experiment 2) post-transplant). *, *P* < 0.05, Mann-Whitney test. **(D)** Uniform manifold approximation and projection (UMAP) plot shows the cell clusters from KPC*^Smad4^*^-WT^ (n=3) or KPC*^Smad4^*^-KO^ (n=6) tumors analyzed by single-cell RNA-sequencing (scRNA-seq). Different genotypes are color coded. **(E)** Upregulated and downregulated differentially expressed genes (DEGs) in each cell type identified by pseudobulk analysis from the scRNA-seq of KPC*^Smad4^*^-WT^ or KPC*^Smad4^*^-KO^ tumors. False discovery rate (FDR) < 0.05. **(F)** Cell type contribution in KPC*^Smad4^*^-WT^ or KPC*^Smad4^*^-KO^ tumors, represented as bar plots showing proportions of the different cell clusters in each condition. **(G)** Flow cytometric analysis of neutrophils (CD45^+^CD11b^+^Gr1^+^) and macrophages (CD45^+^Gr1^-^CD11b^+^F4/80^+^) from live singlets in KPC*^Smad4^*^-WT^ or KPC*^Smad4^*^-KO^ tumors in nu/nu mice. Results show mean ± SEM from 3 separate experiments, each with 1 WT group and 2 groups of KO pools from 2 different guides. **, *P* < 0.01; ***, *P* < 0.001, Mann-Whitney test. **(H)** UMAP plot of neutrophils from KPC*^Smad4^*^-WT^ (n=3) or KPC*^Smad4^*^-KO^ (n=6) tumors analyzed by scRNA-seq. Different sub-clusters are color coded. **(I)** Sub-cluster contribution in neutrophils of KPC*^Smad4^*^-WT^ or KPC*^Smad4^*^-KO^ tumors, represented as bar plots showing proportions of the different sub-clusters in each condition. **(J)** UMAP plot of macrophages from KPC*^Smad4^*^-WT^ (n=3) or KPC*^Smad4^*^-KO^ (n=6) tumors analyzed by scRNA-seq. Different sub-clusters are color coded. **(K)** Sub-cluster contribution in macrophages of KPC*^Smad4^*^-WT^ or KPC*^Smad4^*^-KO^ tumors, represented as bar plots showing proportions of the different sub-clusters in each condition. **(L)** Dot plot visualization of the scaled average expression of macrophage markers in macrophages from KPC*^Smad4^*^-WT^ or KPC*^Smad4^*^-KO^ tumors analyzed by scRNA-seq. The color intensity represents the expression level and the size of the dots represents the percentage of expressing cells. **(M)** Selected significantly upregulated (i.e. NES > 1.50 and FDR < 0.25) and downregulated (i.e. NES < -1.50 and FDR < 0.25) pathways identified by gene set enrichment analysis (GSEA) of macrophages from KPC*^Smad4^*^-KO^ compared to KPC*^Smad4^*^-WT^ tumors, as assessed by pseudobulk analysis from the scRNA-seq dataset. NES, normalized enrichment score. **(N)** Cell-cell communication analysis using CellChat showing the number of connections lost (in blue) or gained (in red) between malignant cells, fibroblasts, macrophages and neutrophils in KPC*^Smad4^*^-KO^ tumors compared to KPC*^Smad4^*^-WT^ tumors, as assessed by scRNA-seq. **(O)** Selected ligand-receptor interactions and their strength based on CellChat analysis between malignant cells, fibroblasts, macrophages and neutrophils in KPC*^Smad4^*^-KO^ tumors compared to KPC*^Smad4^*^-WT^ tumors. **(P-Q)** Selected pathways with significantly different connections between malignant cells, fibroblasts, macrophages, and neutrophils in KPC*^Smad4^*^-KO^ tumors compared to KPC*^Smad4^*^-WT^ tumors.

To understand how SMAD4 loss in KPC malignant cells might impact the TME, we first generated single-cell RNA-sequencing (scRNA-seq) profiles of KPC*^Smad4-^*^WT^ and KPC*^Smad4^*^-KO^ PDAC orthotopic allografts in which the malignant cells were engineered to express green fluorescent protein (GFP; **Fig. 2D**; **Supplementary Fig. S2H-M** and **Supplementary Table S3**). scRNA-seq profiles of malignant cells could be readily distinguished from non-malignant cells by the expression of *Gfp* transcripts and inferred copy number variations (CNV; **Supplementary Fig. S2K-L**). *Smad4* deletion markedly altered the transcriptomes of KPC*^Smad4^*^-KO^ relative to KPC*^Smad4^*^-WT^ malignant cells (6,121 genes differentially expressed, FDR < 0.05). Remarkably, scRNA-seq profiles also revealed significant and selective changes in the transcriptomes and proportions of CAFs, macrophages and neutrophils in KPC*^Smad4^*^-KO^ relative to KPC*^Smad4^*^-WT^ tumors (**Fig. 2E-F**). Thus, SMAD4 loss in PDAC malignant cells impacts CAFs and innate immune cells in the TME.

Flow cytometric analysis confirmed an increase in neutrophils and a decrease in macrophages in KPC*^Smad4^*^-KO^ relative to KPC*^Smad4^*^-WT^ tumors (**Fig. 2G**; **Supplementary Fig. S2N-P**). Analysis of neutrophil clusters within our scRNA-seq profiles identified T1 immature, T2 mature and T3 pro-angiogenic/tumor-promoting neutrophils that have been previously described in PDAC (36), as well as a small subset expressing T1 markers and high levels of *Mpo*, *Ly6g* and *Ly6c1* transcripts (**Fig. 2H-I**; **Supplementary Fig. S2Q**). Of note, markers of T3 neutrophils were upregulated in KPC*^Smad4^*^-KO^ tumors (**Supplementary Fig. S2R**). Analysis of macrophage clusters identified seven sub-clusters similar to recently described types of tumor-associated macrophages (TAMs) (13), as well as a small cluster, which we named Insyn2b-TAMs based on marker expression (**Fig. 2J-K**; **Supplementary Fig. S2S**). Of note, previously described interferon-primed TAMs (IFN-TAMs), lipid-associated TAMs (LA-TAMs) and inflammatory cytokine-enriched TAMs (Inflam-TAMs) appeared more abundant in KPC*^Smad4^*^-KO^ relative to KPC*^Smad4^*^-WT^ PDAC, while proliferating TAMs (Prolif-TAMs) were reduced (**Fig. 2L-M**).

CellChat analysis (37), which infers patterns of cell-cell communication from scRNA-seq profiles, identified potential changes in communication among malignant cells, neutrophils, macrophages and CAFs in KPC*^Smad4^*^-KO^ relative to KPC*^Smad4^*^-WT^ PDAC (**Fig. 2N**). These included a role for malignant cell-derived tumor necrosis factor (TNF) in dictating macrophage composition of KPC*^Smad4^*^-KO^ PDAC, as well as macrophage- and malignant cell-derived chemokine (C-X-C motif) ligand 1 and 2 (CXCL1 and CXCL2) in the recruitment of neutrophils to KPC*^Smad4^*^-KO^ PDAC (**Fig. 2O-Q**). Fibroblasts in KPC*^Smad4^*^-KO^ tumors also appeared to be more involved in neutrophil recruitment via the CXCR2 pathway compared to KPC*^Smad4^*^-WT^ tumors, as previously demonstrated in iCAF-rich PDAC (20) (**Fig. 2O**). Of note, the most affected interaction in KPC*^Smad4^*^-KO^ PDAC appeared to be between malignant cells and fibroblasts, in line with CAFs being the most impacted stromal cell population upon SMAD4 loss (2,294 genes differentially expressed, FDR < 0.05; **Fig. 2N** and **2E**).

These analyses show that loss of SMAD4 in KRAS^G12D^ p53 mutant PDAC malignant cells shapes the immune TME, and suggest that CAFs are also profoundly impacted.

### *Smad4* loss drives a fibro-inflammatory stroma in KPC PDAC

Collagen deposition was significantly reduced in KPC*^Smad4^*^-KO^ tumors relative to KPC*^Smad4^*^-WT^ tumors, further suggesting that malignant cell SMAD4 status impacts the CAF composition in PDAC (**Supplementary Fig. S3A-C**). Indeed, the proportions of iCAFs, myCAFs and apCAFs defined by scRNA-seq profiles were significantly different between KPC*^Smad4^*^-KO^ tumors and KPC*^Smad4^*^-WT^ PDAC (**Fig. 3A-C**; **Supplementary Fig. S3D-E**). Of note, scRNA-seq analysis inferred a decrease in the myCAF/iCAF ratio in KPC*^Smad4^*^-KO^ tumors, which we confirmed by flow cytometry (myCAF, Ly6C^-^/MHCII^-^; iCAF, Ly6C^+^/MHCII^-^; **Fig. 3C-F**; **Supplementary Fig. S3E-G**). In keeping with these findings, gene set enrichment analysis (GSEA) of CAFs showed that iCAF-associated pathways were significantly upregulated in KPC*^Smad4^*^-KO^ tumors (5) (**Fig. 3G**). Moreover, proliferation-associated pathways, which were previously shown to be enriched in myCAFs compared to iCAFs (5), were downregulated (**Fig. 3G**). ApCAF abundance measured by flow cytometry was also lower in KPC*^Smad4^*^-KO^ tumors relative to KPC*^Smad4^*^-WT^ controls (apCAF, Ly6C^-^/MHCII^+^; **Fig. 3D-E**; **Supplementary Fig. S3F**).

**Figure 3.**
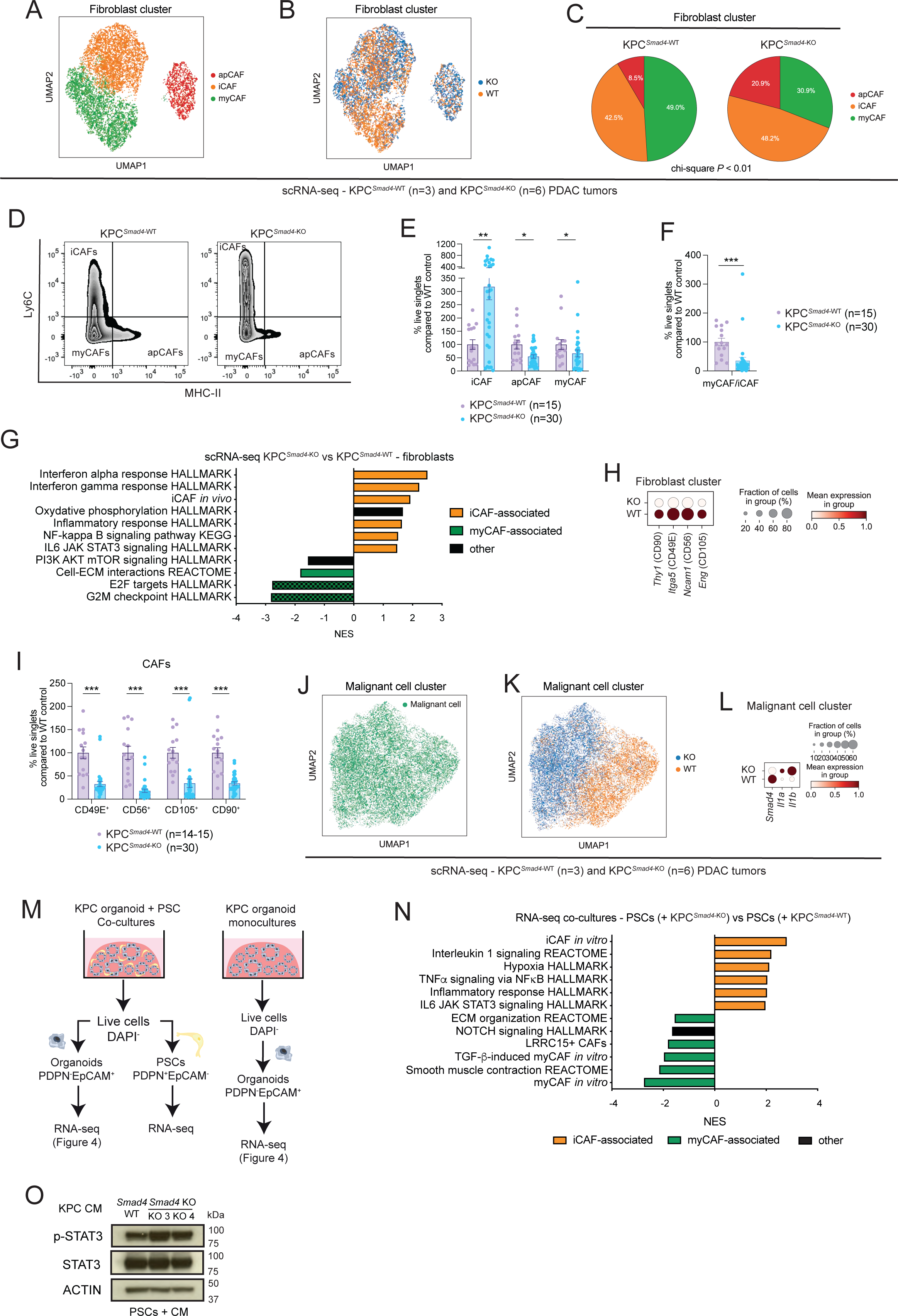
*Smad4* loss drives a fibro-inflammatory stroma in KPC PDAC. **(A-B)** UMAP plots showing the cell cluster of cancer-associated fibroblasts (CAFs) from KPC*^Smad4^*^-WT^ (n=3) or KPC*^Smad4^*^-KO^ (n=6) PDAC tumors analyzed by scRNA-seq. Different CAF clusters **(A)** or genotypes **(B)** are color coded. **(C)** Pie charts showing proportions of different CAF clusters from KPC*^Smad4^*^-WT^ or KPC*^Smad4^*^-KO^ tumors. *P*, chi-square test. **(D)** Representative flow plots of Ly6C^-^ MHCII^-^ myCAFs, Ly6C^+^MHCII^-^ iCAFs and Ly6C^-^MHCII^+^ apCAFs from KPC*^Smad4^*^-WT^ or KPC*^Smad4^*^-KO^ tumors in C57BL/6J mice. **(E)** Flow cytometric analyses of myCAFs (Ly6C^-^MHCII^-^), iCAFs (Ly6C^+^MHCII^-^) and apCAFs (Ly6C^-^MHCII^+^) from live singlets in KPC*^Smad4^*^-WT^ or KPC*^Smad4^*^-KO^ tumors in C57BL/6J mice. Results show mean ± SEM from 3 separate experiments, each with 1 WT group and 2 groups of KO pools from 2 different guides. *, *P* < 0.05, **\***\*, *P* < 0.01, Mann-Whitney test. **(F)** Flow cytometric analyses of myCAF/iCAF ratio from live singlets in KPC*^Smad4^*^-WT^ or KPC*^Smad4^*^-KO^ tumors in C57BL/6J mice. Results show mean ± SEM from 3 separate experiments, each with 1 WT group and 2 groups of KO pools from 2 different guides. ***, *P* < 0.001, Mann-Whitney test. **(G)** Selected significantly upregulated (i.e. NES > 1.50 and FDR < 0.25; apart for the IL6 JAK STAT3 signaling HALLMARK which has NES=1.47) and downregulated (i.e. NES < -1.50 and FDR < 0.25) pathways identified by GSEA of CAFs from KPC*^Smad4^*^-KO^ compared to KPC*^Smad4^*^-WT^ tumors, as assessed by pseudobulk analysis from the scRNA-seq dataset. The *in vivo* iCAF signature is from Elyada et al (3). **(H)** Dot plot visualization of the scaled average expression of myCAF-enriched markers in CAFs from KPC*^Smad4^*^-WT^ or KPC*^Smad4^*^-KO^ tumors, as analyzed by scRNA-seq. The color intensity represents the expression level and the size of the dots represents the percentage of expressing cells. **(I)** Flow cytometric analyses of CD90^+^, CD49E^+^, CD56^+^ and CD105^+^ CAFs from live singlets in KPC*^Smad4^*^-WT^ or KPC*^Smad4^*^-KO^ tumors in C57BL/6J mice. Results show mean ± SEM from 3 separate experiments, each with 1 WT group and 2 groups of KO pools from 2 different guides. ***, *P* < 0.001, Mann-Whitney test. **(J-K)** UMAP plots showing the malignant cell cluster from KPC*^Smad4^*^-WT^ (n=3) or KPC*^Smad4^*^-KO^ (n=6) tumors analyzed by scRNA-seq **(J)**. Different genotypes are color coded **(K)**. **(L)** Dot plot visualization of the scaled average expression of *Smad4*, *Il1a* and *Il1b* in malignant cells from KPC*^Smad4^*^-WT^ or KPC*^Smad4^*^-KO^ tumors, as analyzed by scRNA-seq. The color intensity represents the expression level and the size of the dots represents the percentage of expressing cells. **(M)** Schematic of flow-sorting strategy of pancreatic stellate cells (PSCs) and malignant cells from monocultures or co-cultures of KPC*^Smad4^*^-WT^ or KPC*^Smad4^*^-KO^ PDAC organoids for RNA-sequencing (RNA-seq) analysis. **(N)** Selected significantly upregulated (i.e. NES > 1.50 and FDR < 0.25) and downregulated (i.e. NES < -1.50 and FDR < 0.25) pathways identified by GSEA of PSCs cultured with KPC*^Smad4^*^-KO^ organoids (n=10) compared to PSCs cultured with KPC*^Smad4^*^-WT^ organoids (n=5). The *in vitro* iCAF and myCAF signatures are from Öhlund et al (4). The TGF-β-induced myCAF *in vitro* signature is from Mucciolo and Araos Henríquez et al (11). The LRRC15^+^ CAF signature was obtained from Dominguez et al (7), as published in Mucciolo and Araos Henríquez et al (11). **(O)** Western blot analysis of phospho-STAT3 (p-STAT3) and STAT3 in murine PSCs cultured for 4 days in PDAC organoid conditioned media (CM) from KPC*^Smad4^*^-WT^ or KPC*^Smad4^*^-KO^ organoids. ACTIN, loading control.

Since we previously identified heterogeneity among PDAC myCAFs (11), we further interrogated these CAFs by flow cytometry using CD90 (encoded by *Thy1*), which marks a subset of PDAC myCAFs (11), as well as CD105 (encoded by *Eng*, and a CAF-lineage marker (6)), CD49E (encoded by *Itga5*) and CD56 (encoded by *Ncam1*). CD105, CD49E and CD56 have not been described previously as myCAF markers but were enriched in myCAFs by scRNA-seq (**Supplementary Fig. S3D**). Each of these myCAF markers was significantly downregulated in CAFs in KPC*^Smad4^*^-KO^ tumors relative to KPC*^Smad4^*^-WT^ tumors, while tumor-promoting CD90^-^ myCAFs were increased (11) (**Fig. 3H-I**; **Supplementary Fig. S3H-J**). Furthermore, malignant cell scRNA-seq profiles in KPC*^Smad^*^-KO^ tumors expressed high levels of *Il1a* and *Il1b*, which direct iCAF formation (5) (**Fig. 3J-L**). Thus, loss of SMAD4 in KPC malignant cells profoundly shapes CAF composition in PDAC, potentially through interleukin 1 (IL-1) malignant cell-CAF signaling.

To further deconvolute changes in malignant cell-fibroblast crosstalk upon SMAD4 loss, we leveraged our validated co-culture model of PDAC organoids and pancreatic stellate cells (PSCs). PSCs are precursors of CAFs and can model iCAFs and myCAFs *in vitro* (4,5,9,11). KPC*^Smad4^*^-KO^ and KPC*^Smad4^*^-WT^ organoids cultured either alone or together with PSCs were flow-sorted and analyzed by RNA-seq (**Fig. 3M**; **Supplementary Fig. S3K** and **Supplementary Tables S4-S5**). In keeping with our *in vivo* observations, iCAF markers and associated pathways were significantly increased, while myCAF markers and associated pathways were decreased in KPC*^Smad4^*^-KO^/PSC relative to KPC*^Smad4^*^-WT^/PSC co-cultures (**Fig. 3N**; **Supplementary Fig. S3L**). Furthermore, the JAK/STAT pathway which is required to maintain the iCAF phenotype (5), was enriched in PSCs co-cultured with KPC*^Smad4^*^-KO^ organoids (**Fig. 3N**). Phospho-STAT3 (p-STAT3) levels confirmed enhanced JAK/STAT activation in PSCs cultured with conditioned media (CM) from KPC*^Smad4^*^-KO^ relative to KPC*^Smad4^*^-WT^ organoids (**Fig. 3O**).

These analyses demonstrate that *Smad4* deletion in KRAS^G12D^ p53 mutant PDAC malignant cells directly shapes CAF composition towards a more inflammatory phenotype.

### *Smad4* loss upregulates IL-1 and JAK/STAT signaling in KPC PDAC

To identify mediators of malignant cell-fibroblast crosstalk in *Smad4*-deleted PDAC, we evaluated bidirectional signaling between malignant cells and CAFs in PDAC organoid/PSC co-cultures. Ligand-receptor interaction analysis using NicheNet pinpointed *Il1a* as the top malignant cell-produced mediator of the malignant-CAF crosstalk in KPC*^Smad4^*^-KO^/PSC relative to KPC*^Smad4^*^-WT^/PSC co-cultures (**Fig. 4A**; **Supplementary Fig. S4A** and **Supplementary Table S6**) (38). Indeed, *Il1a* expression was significantly upregulated in KPC*^Smad4^*^-KO^ organoid monocultures compared to KPC*^Smad4^*^-WT^ controls (**Fig. 4B**). Since we showed previously that IL-1 signaling is a key driver of the iCAF phenotype (5), these data support a model in which increased IL-1 expression following *Smad4* loss from malignant cells enhances iCAF generation. Of note, *Il1a* levels in organoids were further upregulated in KPC*^Smad4^*^-KO^/PSC co-cultures compared to monoculture, suggesting a positive feedback loop in the presence of an iCAF-rich environment (**Fig. 4B**). Indeed, CAF-produced *Il1a* also appeared to be the top mediator of both CAF-CAF crosstalk and CAF-malignant cell crosstalk in KPC*^Smad4^*^-KO^/PSC co-cultures (**Fig. 4C-D**; **Supplementary Fig. S4B-C**). Furthermore, *Tnf* scored as the top mediator of malignant cell-malignant cell crosstalk in KPC*^Smad4^*^-KO^/PSC co-cultures (**Supplementary Fig. S4D-E**). In line with this, *Tnf* expression was upregulated upon SMAD4 loss in the malignant cells both *in vitro* and *in vivo*, but it was not further upregulated when the organoids were co-cultured with PSCs (**Supplementary Fig. S4F-G**).

**Figure 4.**
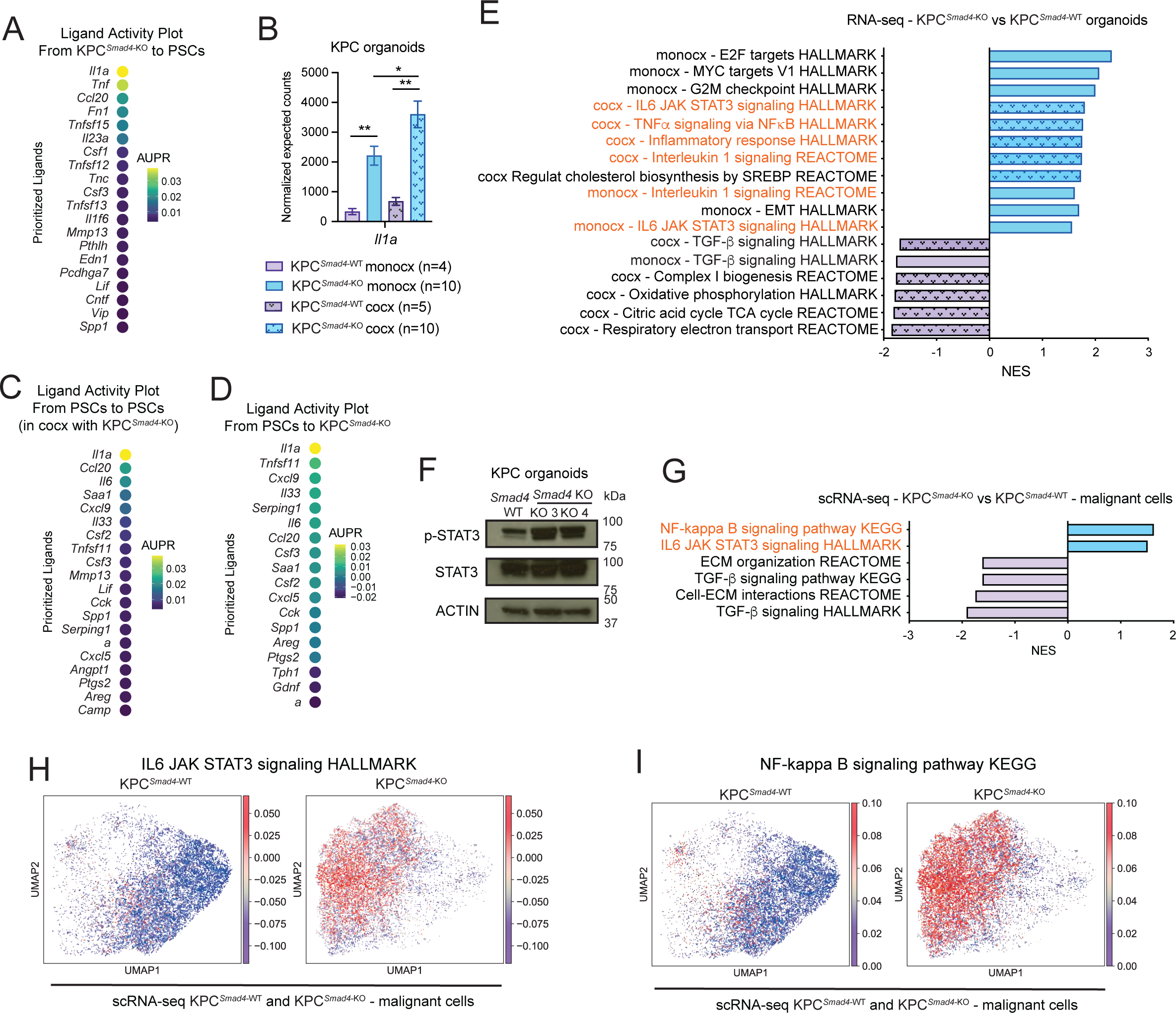
*Smad4* loss upregulates IL-1 and JAK/STAT signaling in KPC PDAC. **(A)** Ligand activity plot shows the top ligands of KPC*^Smad4^*^-KO^ PDAC organoids that regulate target genes in co-cultured PSCs, as assessed by NicheNet analysis of RNA-seq. AUPR, area under the precision-recall curve. **(B)** RNA-seq expression of *Il1a* in KPC*^Smad4^*^-WT^ or KPC*^Smad4^*^-KO^ organoids flow-sorted from monocultures or co-cultures with PSCs. Results show mean ± SEM. *, *P* < 0.05, **, *P* < 0.01, Mann-Whitney test. **(C)** Ligand activity plot shows the top ligands of PSCs in co-culture with KPC*^Smad4^*^-KO^ organoids that regulate target genes in PSCs, as assessed by NicheNet analysis of RNA-seq. **(D)** Ligand activity plot shows the top ligands from PSCs that regulate target genes in co-cultured KPC*^Smad4^*^-KO^ organoids, as assessed by NicheNet analysis of RNA-seq. **(E)** Selected pathways found significantly enriched or depleted (NES > 1.5 or < -1.5; FDR < 0.25) by GSEA in KPC*^Smad4^*^-KO^ malignant cells (n=10 co-cultures, n=10 monocultures) compared to KPC*^Smad4^*^-WT^ malignant cells (n=5 co-cultures, n=4 monocultures) flow-sorted from monocultures or co-cultures with PSCs. The smooth pattern highlights the GSEA of the monocultures. Inflammatory pathways are highlighted in orange. Cocx, co-culture; monocx, monoculture. **(F)** Western blot analysis of phospho-STAT3 (p-STAT3) and STAT3 in KPC*^Smad4^*^-WT^ or KPC*^Smad4^*^-KO^ organoids cultured in reduced media. ACTIN, loading control. **(G)** Selected significantly upregulated (i.e. NES > 1.50 and FDR < 0.05) or significantly downregulated (i.e. NES < -1.50 and FDR < 0.05) pathways identified by GSEA of malignant cells from KPC*^Smad4^*^-KO^ tumors (n=6) compared to malignant cells from KPC*^Smad4^*^-WT^ tumors (n=3), as assessed by pseudobulk analysis of scRNA-seq. Inflammatory pathways are highlighted in orange. **(H-I)** UMAP plots of malignant cells from KPC*^Smad4^*^-WT^ or KPC*^Smad4^*^-KO^ tumors colored by the normalized expression score of the HALLMARK IL6 JAK STAT3 signaling **(H)** or the KEGG NF-κB signaling **(I)** pathways.

In line with these findings, IL-1 and NF-κB signaling were enriched in KPC*^Smad4^*^-KO^ relative to KPC*^Smad4^*^-WT^ monocultures and further enhanced when co-cultured with PSCs (**Fig. 4E**). Interestingly, JAK/STAT signaling and p-STAT3 levels were also upregulated in KPC*^Smad4^*^-KO^ relative to KPC*^Smad4^*^-WT^ monocultures, and the JAK/STAT pathway was further enriched when co-cultured with PSCs (**Fig. 4E-F**). Finally, while TGF-β signaling was downregulated, gene expression signatures associated with NF-κB and JAK/STAT signaling were also upregulated in scRNA-seq profiles of KPC*^Smad4^*^-KO^ malignant cells *in vivo* (**Fig. 4G-I**; **Supplementary Fig. S4H**).

Our analyses suggest that *Smad4* loss from KRAS^G12D^ p53 mutant PDAC malignant cells upregulates IL-1 expression to promote iCAF formation and enhances JAK/STAT signaling activation.

### *Smad4* loss tunes TME crosstalk and signaling dependencies in PDAC with distinct KRAS status

The *KRAS*^G12V^ mutant allele also occurs frequently in human PDAC but is rarely studied in pre-clinical models (25). Therefore, we assessed whether *Smad4* loss would impact KRAS^G12V^ mutant PDAC similarly to that observed in KRAS^G12D^ mutant PDAC. To do this, we used PDAC organoids generated from the *Kras^FRT-LSL-^*^G12V*-FRT/+*^; *Trp53^LSL-^*^R172H^; *Pdx1-Cre*; *Rosa26-FlpO^ERT2^* (hereafter, KvPC) mouse model (39). PDAC formation in this GEMM is driven by the same *Trp53* mutant allele and *Cre* promoter as the KPC model, but differs by having *Kras*^G12V^ instead of *Kras*^G12D^ (26,39). We generated isogenic *Smad4* KO (KvPC*^Smad4^*^-KO^) and *Smad4* WT (KvPC*^Smad4^*^-WT^) KvPC organoids exactly as for KPC organoids (34) (**Supplementary Fig. S5A-D**).

As observed for KPC organoid-derived mouse models, allografted KvPC*^Smad4^*^-KO^ organoids formed tumors that grew significantly faster than KvPC*^Smad4^*^-WT^ PDAC (**Fig. 5A-B**; **Supplementary Fig. S5E**). Like KPC*^Smad4^*^-KO^ PDAC, KvPC*^Smad4^*^-KO^ PDAC also contained more neutrophils and fewer macrophages than KvPC*^Smad4^*^-WT^ PDAC (**Fig. 5C-E**; **Supplementary Fig. S5F-J** and **Supplementary Table S7**). Furthermore, certain malignant cell-stromal interactions were similarly altered in both KRAS mutant-driven models upon SMAD4 loss, including the potential role of *Smad4* KO malignant cell-produced TNF in shaping macrophage composition and macrophage-produced CXCL1 and CXCL2 in recruiting neutrophils in KvPC*^Smad4^*^-KO^ PDAC (**Fig. 5F-H**; **Supplementary Fig. S5K**).

**Figure 5.**
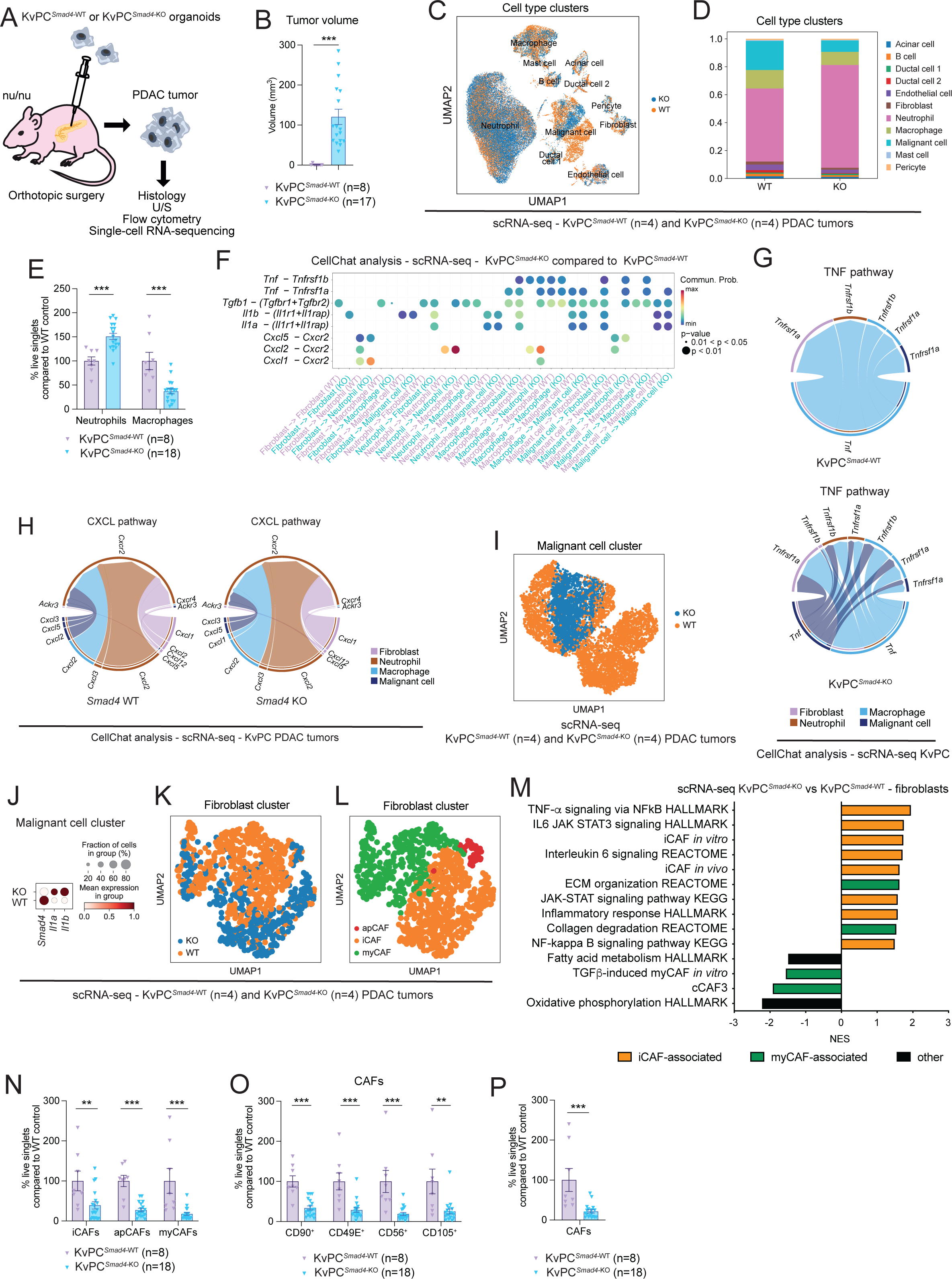
*Smad4* loss drives a fibro-inflammatory stroma in KvPC PDAC. **(A)** Schematic of analyses of KvPC organoid-derived PDAC models in nu/nu mice. **(B)** Tumor volumes as measured by ultrasound-based imaging of tumors derived from the transplantation of *Smad4* WT or *Smad4* KO KvPC (i.e. *Kras*^G12V^ p53 mutant) PDAC organoids. Results show mean ± SEM from 2 separate experiments, each with 1 WT group and 2 groups of KO pools from 2 different guides (n=8-17 mice/cohort; at day 29 (experiment 1) or day 30 (experiment 2) post-transplant). ***, *P* < 0.001, Mann-Whitney test. **(C)** UMAP plot of all cell types from KvPC*^Smad4^*^-WT^ (n=4) or KvPC*^Smad4^*^-KO^ (n=4) PDAC tumors analyzed by scRNA-seq. Different genotypes are color coded. **(D)** Cell type contribution in KvPC*^Smad4^*^-WT^ or KvPC*^Smad4^*^-KO^ tumors, represented as bar plots showing proportions of the different cell clusters in each condition. **(E)** Flow cytometric analysis of neutrophils (CD45^+^CD11b^+^Gr1^+^) and macrophages (CD45^+^Gr1^-^CD11b^+^F4/80^+^) from live singlets in KvPC*^Smad4^*^-WT^ or KvPC*^Smad4^*^-KO^ tumors. Results show mean ± SEM from 2 separate experiments, each with 1 WT group and 2 groups of KO pools from 2 different guides. ***, *P* < 0.001, Mann-Whitney test. **(F)** Selected ligand-receptor interactions and their strength based on CellChat analysis between malignant cells, fibroblasts, macrophages, and neutrophils in KvPC*^Smad4^*^-KO^ tumors compared to KvPC*^Smad4^*^-WT^ tumors. **(G-H)** Selected pathway with significantly different connections between malignant cells, fibroblasts, macrophages, and neutrophils in KvPC*^Smad4^*^-KO^ tumors compared to KvPC*^Smad4^*^-WT^ tumors. **(I)** UMAP plot of malignant cells from KvPC*^Smad4^*^-WT^ (n=4) or KvPC*^Smad4^*^-KO^ (n=4) tumors analyzed by scRNA-seq. Different genotypes are color coded. **(J)** Dot plot visualization of the scaled average expression of *Smad4*, *Il1a* and *Il1b* in malignant cells of KvPC*^Smad4^*^-WT^ or KvPC*^Smad4^*^-KO^ tumors, as analyzed by scRNA-seq. The color intensity represents the expression level and the size of the dots represents the percentage of expressing cells. **(K-L)** UMAP plots of CAFs from KvPC*^Smad4^*^-WT^ (n=4) or KvPC*^Smad4^*^-KO^ (n=4) tumors analyzed by scRNA-seq. Different genotypes **(K)** or CAF sub-clusters **(L)** are color coded. **(M)** Selected significantly upregulated (i.e. NES > 1.50 and FDR < 0.25; apart from the NF-kappa B signaling pathway, which has NES = 1.49) and downregulated (i.e. NES < -1.50 and FDR < 0.25; apart from the fatty acid metabolism, which has NES = -1.48) pathways identified by GSEA of CAFs from KvPC*^Smad4^*^-KO^ compared to KvPC*^Smad4^*^-WT^ tumors, as assessed by pseudobulk analysis from the scRNA-seq dataset. The *in vivo* iCAF signature is from Elyada et al (3). The *in vitro* iCAF signature is from Öhlund et al (4). The TGF-β-induced myCAF *in vitro* signature is from Mucciolo and Araos Henríquez et al (11). The cCAF3 signature was obtained from McAndrews et al (8), as published in Mucciolo and Araos Henríquez et al (11). **(N)** Flow cytometric analysis of myCAFs (Ly6C^-^MHCII^-^), iCAFs (Ly6C^+^MHCII^-^) and apCAFs (Ly6C^-^MHCII^+^) from live singlets in KvPC*^Smad4^*^-WT^ or KvPC*^Smad4^*^-KO^ tumors. Results show mean ± SEM from 2 separate experiments, each with 1 WT group and 2 groups of KO pools from 2 different guides. **, *P* < 0.01; ***, *P* < 0.001, Mann-Whitney test. **(O)** Flow cytometric analyses of CD90^+^, CD49E^+^, CD56^+^ and CD105^+^ CAFs from live singlets in KvPC*^Smad4^*^-WT^ or KvPC*^Smad4^*^-KO^ tumors. Results show mean ± SEM from 2 separate experiments, each with 1 WT group and 2 groups of KO pools from 2 different guides. **, *P* < 0.01; ***, *P* < 0.001, Mann-Whitney test. **(P)** Flow cytometric analysis of CAFs (CD45^-^CD31^-^EpCAM^-^PDPN^+^) from live singlets in KvPC*^Smad4^*^-WT^ or KvPC*^Smad4^*^-KO^ tumors. Results show mean ± SEM from 2 separate experiments, each with 1 WT group and 2 groups of KO pools from 2 different guides. ***, *P* < 0.001, Mann-Whitney test.

Moreover, as observed in KPC*^Smad4^*^-KO^ PDAC, *Il1a* and *Il1b* expression were also upregulated in KvPC*^Smad4^*^-KO^ PDAC malignant cells compared to KvPC*^Smad4^*^-WT^ tumors (**Fig. 5I-J**). Furthermore, *Smad4* loss from KvPC malignant cells produced PDAC with significantly less collagen deposition compared to WT controls (**Supplementary Fig. S5L-N**). In line with these observations, scRNA-seq profiles of KvPC*^Smad4^*^-KO^ PDAC-derived CAFs revealed a decrease in myCAF-associated pathways, while iCAF-associated pathways were significantly increased (**Fig. 5K-M**; **Supplementary Fig. S5O-P**). Moreover, flow cytometry analysis confirmed loss of myCAFs, apCAFs and CD90^+^, CD49E^+^, CD56^+^ and CD105^+^ myCAF populations in KvPC*^Smad4^*^-KO^ PDAC tumors compared to WT controls, as observed in KPC tumors (**Fig. 5N-O**; **Supplementary Fig. S5Q**).

Despite these similarities, the myCAF/iCAF ratio was not significantly downregulated by flow cytometric analysis in KvPC*^Smad4^*^-KO^ PDAC tumors compared to WT controls, and iCAF abundance was also reduced, likely because of total CAF abundance downregulation in KvPC*^Smad4^*^-KO^ PDAC (**Fig. 5N** and **5P**; **Supplementary Fig. S5R**). Moreover, while the same four neutrophil states were identified in both KPC and KvPC tumors, their patterns of marker expression were in part differentially altered following SMAD4 loss (**Supplementary Fig. S6A-D**). Furthermore, while Inflam-TAMs also appeared more abundant in KvPC*^Smad4^*^-KO^ PDAC, as observed in KPC*^Smad4^*^-KO^ tumors, Prolif-TAMs were not clearly decreased, and markers of IFN-TAMs (not found as a defined sub-cluster in KvPC tumors) or LA-TAMs were not clearly increased in KvPC*^Smad4^*^-KO^ PDAC compared to WT controls (**Supplementary Fig. S6E-I**). These data suggest that SMAD4 loss has different effects on the stroma composition of PDAC tumors with distinct KRAS mutations.

Most strikingly, the JAK/STAT and NF-κB signaling pathways were not upregulated in KvPC*^Smad4^*^-KO^ malignant cells compared to KvPC*^Smad4^*^-WT^ PDAC *in vivo*, as analyzed by scRNA-seq (**Fig. 6A**). Thus, to further explore how SMAD4 loss directly impacts KvPC PDAC malignant cells, we established co-cultures of PSCs and KvPC PDAC organoids and analyzed both flow-sorted populations by RNA-seq, as done for KPC organoids (**Fig. 6B**; **Supplementary Fig. S6J** and **Supplementary Tables S8-S9**). Validating our *in vivo* observations, this analysis showed that iCAF markers and associated pathways were upregulated, and myCAF-associated pathways were downregulated, when PSCs were co-cultured with KvPC*^Smad4^*^-KO^ organoids, albeit with some differences compared to KPC*^Smad4^*^-KO^ co-cultures (**Fig. 6C**; **Supplementary Fig. S6K**). Of note, as observed *in vivo*, GSEA of KvPC malignant cells showed that JAK/STAT signaling was not upregulated following SMAD4 loss (**Fig. 6D**). This was confirmed by protein analysis of p-STAT3 levels, which were not upregulated in KvPC organoids following SMAD4 loss (**Fig. 6E**).

**Figure 6.**
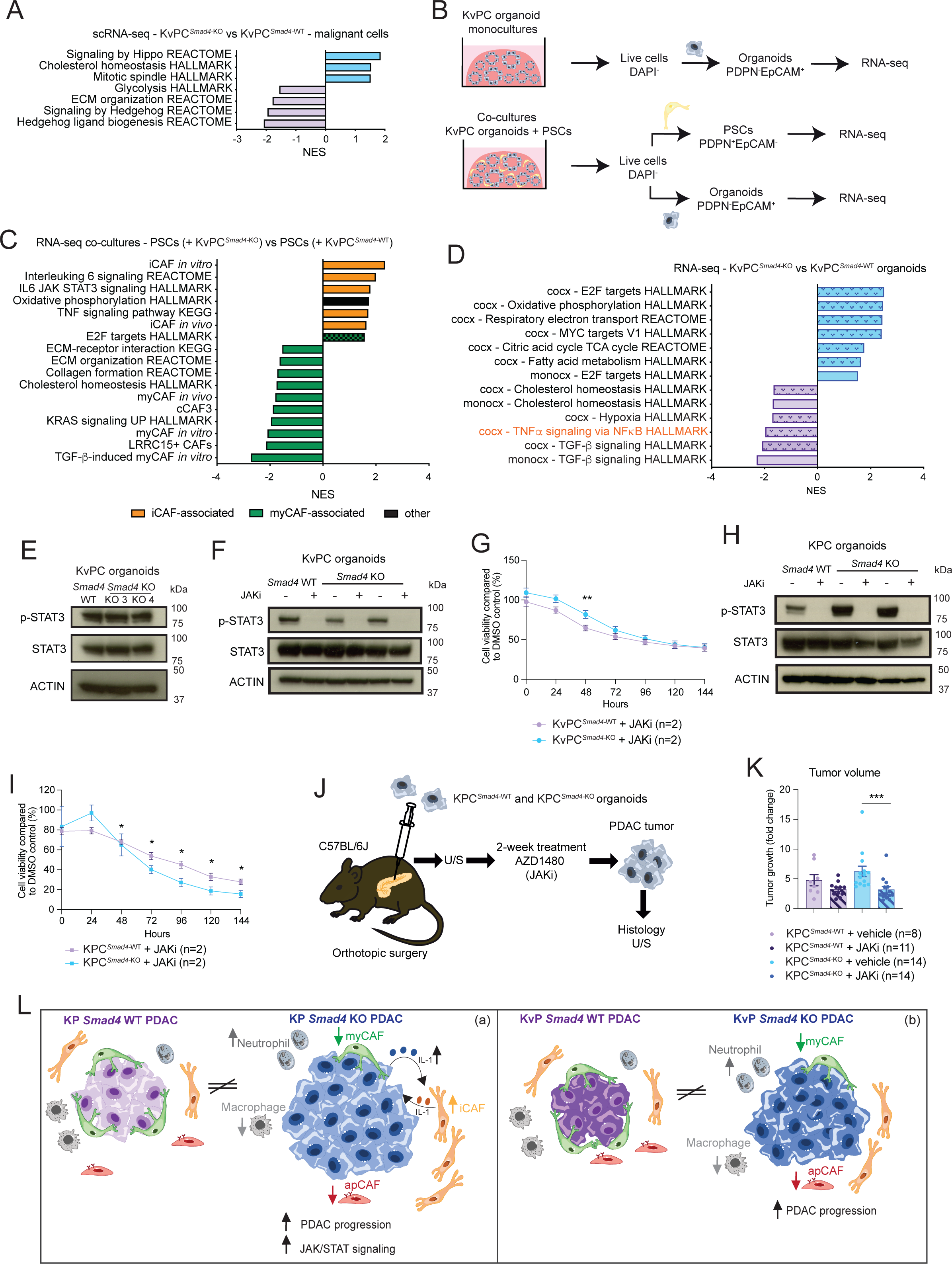
*Smad4* loss tunes signaling dependencies in PDAC with distinct KRAS status. **(A)** Selected significantly upregulated (i.e NES > 1.50 and FDR < 0.25) or significantly downregulated (i.e NES < -1.50 and FDR < 0.25) pathways identified by GSEA of malignant cells from KvPC*^Smad4^*^-KO^ PDAC tumors (n=4) compared to malignant cells from KvPC*^Smad4^*^-WT^ tumors (n=4), as assessed by pseudobulk analysis of the scRNA-seq dataset. **(B)** Schematic of flow-sorting strategy of PSCs and malignant cells from monocultures or co-cultures of KvPC*^Smad4^*^-WT^ or KvPC*^Smad4^*^-KO^ PDAC organoids for RNA-seq analysis. **(C)** Selected significantly upregulated (i.e. NES > 1.50 and FDR < 0.25) and downregulated (i.e. NES < -1.50 and FDR < 0.25) pathways identified by GSEA of PSCs cultured with KvPC*^Smad4^*^-KO^ organoids (n=8) compared to PSCs cultured with KvPC*^Smad4^*^-WT^ organoids (n=4). The *in vivo* iCAF and myCAF signatures are from Elyada et al (3). The *in vitro* iCAF and myCAF signatures are from Öhlund et al (4). The TGF-β-induced myCAF *in vitro* signature is from Mucciolo and Araos Henríquez et al (11). The LRRC15^+^ CAF and cCAF3 signatures were obtained from Dominguez et al (7) and McAndrews et al (8), respectively, as published in Mucciolo and Araos Henríquez et al (11). **(D)** Selected pathways found significantly enriched or depleted (NES > 1.5 or < -1.5; FDR < 0.25) by GSEA in KvPC*^Smad4^*^-KO^ malignant cells (n=8 co-cultures, n=7 monocultures) compared to KvPC*^Smad4^*^-WT^ malignant cells (n=4 co-cultures, n=4 monocultures) flow-sorted from monocultures or co-cultures with PSCs. The smooth pattern highlights the GSEA of the monocultures. Inflammatory pathways are highlighted in orange. Cocx, co-culture; monocx, monoculture. **(E)** Western blot analysis of p-STAT3 and STAT3 in KvPC*^Smad4^*^-WT^ or KvPC*^Smad4^*^-KO^ organoids cultured in reduced media. ACTIN, loading control. **(F)** Western blot analysis of p-STAT3 and STAT3 in KvPC*^Smad4^*^-WT^ or KvPC*^Smad4^*^-KO^ organoids cultured for 48 hours in reduced media with or without 8 μM of the JAK inhibitor (JAKi, AZD1480). ACTIN, loading control. **(G)** Proliferation curves of KvPC*^Smad4^*^-WT^ and KvPC*^Smad4^*^-KO^ organoids cultured for 144 hours in Matrigel in reduced media with or without 8 μM JAKi. Data were normalized to the first measurement (at 3 hours post-plating on day 0) and the DMSO control. Results show mean ± SEM of n=2 biological replicates (with n=6 technical replicates each). **, *P* < 0.01, Mann Whitney test. **(H)** Western blot analysis of p-STAT3 and STAT3 in KPC*^Smad4^*^-WT^ or KPC*^Smad4^*^-KO^ organoids cultured for 48 hours in reduced media with or without 8 μM of the JAKi. ACTIN, loading control. **(I)** Proliferation curves of KPC*^Smad4^*^-WT^ and KPC*^Smad4^*^-KO^ organoids cultured for 144 hours in Matrigel in reduced media with or without 8 μM JAKi. Data were normalized to the first measurement (at 3 hours post-plating on day 0) and the DMSO control. Results show mean ± SEM of n=2 biological replicates (with n=6 technical replicates each). *, *P* < 0.05, Mann-Whitney test. **(J)** Schematic of experimental design and downstream analyses of a 2-week treatment study with the JAKi of KPC*^Smad4^*^-WT^ and KPC*^Smad4^*^-KO^ organoid-derived PDAC models in C57BL/6J mice. U/S, ultrasound-based imaging. **(K)** Tumor growth as measured by ultrasound-based imaging of 2-week vehicle- and JAKi-treated KPC*^Smad4^*^-WT^ or KPC*^Smad4^*^-KO^ PDAC tumors. Results show mean ± SEM from 3 separate experiments (n=8-14/cohort). ***, *P* < 0.001, Mann-Whitney test. **(L)** Model illustrating how *Smad4* loss differently shapes murine KRAS^G12D^ (a) and KRAS^G12V^ (b) p53 mutant PDAC tumors.

Since JAK/STAT signaling was upregulated upon *Smad4* deletion in KPC, but not in KvPC, malignant cells, we evaluated whether inhibition of this pathway led to an increased therapeutic sensitivity only in KPC*^Smad4^*^-KO^ PDAC. No significant difference was observed in the proliferation of KvPC*^Smad4^*^-KO^ organoids compared to KvPC*^Smad4^*^-WT^ controls when exposed to the JAK inhibitor (JAKi) AZD1480 (5,40) (**Fig. 6F-G**). In contrast, KPC*^Smad^*^-KO^ organoids were significantly more sensitive to JAK inhibition than KPC*^Smad4^*^-WT^ controls (**Fig. 6H-I**). To test if this difference in JAK signaling dependency might also exist *in vivo*, we treated mice harboring KPC*^Smad^*^-KO^ or KPC*^Smad^*^-WT^ PDACs for two weeks with the JAKi. Remarkably, JAK inhibition significantly impaired the growth of KPC*^Smad^*^-KO^ but not KPC*^Smad^*^-WT^ tumors (**Fig. 6J-K**; **Supplementary Fig. S6L-M**). Thus, distinct mutational patterns in PDAC also impact therapeutic vulnerabilities.

Together, our analyses suggest that SMAD4 loss differently impacts malignant-stromal crosstalk and therapeutic sensitivities in PDAC with distinct KRAS status.

## DISCUSSION

Cancers are rarely driven by single mutations. Distinct combinations of mutations in each tumor can have both malignant cell intrinsic and extrinsic effects. Understanding how different combinations of mutations cooperate to drive the malignant phenotype is key if we are to develop more effective therapies. Here, we show how SMAD4 loss in PDAC malignant cells shapes tumor biology differently in the presence of two distinct KRAS mutations (**Fig. 6L**). While loss of SMAD4 generated a fibro-inflammatory TME in both KPC and KvPC PDAC, it also led to distinct differences in CAFs and innate immune cells in these two tumor types. SMAD4 loss was also associated with an increase in JAK/STAT dependency in KPC*^Smad^*^-KO^, but not KvPC*^Smad^*^-KO^, PDAC compared to *Smad4* WT controls. These mutation-dependent differences in JAK/STAT dependency may explain why JAK inhibitors showed promise in pre-clinical studies but failed to display benefit when used to treat patients with genetically heterogeneous PDAC in clinical trials (5,41–43). Our data suggest that better understanding of how PDAC genetics impact malignant-stromal crosstalk heterogeneity is needed if we are to deploy more effective targeted therapies in the clinic.

Our study also highlights the power of comparing isogenic organoid-derived models to begin to deconvolute the complexity of human PDAC and pinpoint the impact of combinations of mutations on malignant cells and the TME. Further modeling to gradually reconstruct the genetic complexity observed in patients will enable us to deconvolute it further to identify malignant-stromal signatures and therapeutic vulnerabilities common to specific groups of patients. Due to the heterogeneous nature of the TME, scRNA-seq studies coupled with genetic information will be key in future efforts to understand the biology of distinct groups of PDAC and target this more effectively.

Upregulation of STAT3 signaling in the epithelial compartment of KPC GEMMs was also observed following TGFBR2 loss and was associated with increased tissue tension and collagen fiber changes (30,44,45). Here, we found increased STAT3 activation in both malignant cells and the iCAF-rich microenvironment associated with KPC*^Smad^*^-KO^, but not KvPC*^Smad^*^-KO^, PDAC tumors compared to *Smad4* WT controls. While the impact of TGFBR1/2 signaling inhibition in *Smad4* WT and *Smad4* KO PDAC organoids remains to be assessed, our data indicate differences between SMAD4 deletion and TGFBR2 loss. Specifically, contrary to TGFBR2 loss (46), SMAD4 deletion in both KPC and KvPC PDAC drove primary tumor growth *in vivo*. Accordingly, it was previously described that among mutations associated with impaired TGF-β signaling, only SMAD4 loss, but not TGFBR2 mutation, was associated with worse survival of PDAC patients (45). Evidence suggests that inflammatory CAFs play tumor-promoting and immunosuppressive roles in PDAC and other cancer types (2,5,47–49), but whether their increase in *Smad4*-deleted PDAC contributes to this aggressive tumor phenotype remains to be assessed. Moreover, the observed increase in neutrophils may also contribute to the faster growth phenotype of *Smad4* KO KPC and KvPC tumors compared to *Smad4* WT controls. Indeed, therapies targeting neutrophils in pre-clinical mouse models of PDAC have been shown to hinder tumor progression (50).

As KRAS mutant-specific and pan-KRAS inhibitors are being developed (51–54), understanding differences between KRAS mutations, and their associated TMEs, could be key to design effective combinatorial strategies. While our analyses revealed similar effects caused by SMAD4 loss in the context of KPC and KvPC tumors, they also highlighted differences in malignant cell features and stroma composition. Whether such differences are at least partially dependent on intrinsic differences of distinct KRAS mutations remains to be determined. Similarly, it remains to be assessed whether other common mutations in PDAC, such as *CDKN2A* loss, also shape the TME and are required to further stratify *SMAD4*-deficient and *SMAD4* WT tumors.

## MATERIALS AND METHODS

### Human PDAC tissues

Human PDAC tissues used in this study were obtained from surgical resections of patients (both males and females) treated at the University and Hospital Trust of Verona (Azienda Ospedaliera Universitaria Integrata, AOUI) for curative intent. Written informed consent was acquired from patients before specimens’ acquisition. Specimens were acquired under protocols approved by the AOUI Ethics Committee (Comitato Etico Azienda Ospedaliera Universitaria Integrata): approval number 1911 (Prot. n 61413, Prog 1911 on 19/09/2018) and approval number 3456 (Prot. n. 55859, Prog 3456 on 22/09/2021). Clinical information of the human PDAC tissues used, including sex, age, histological diagnosis, staging and genotype information for *KRAS*, *TP53*, *CDKN2A*, *SMAD4* and *TGFBR2*, is provided in **Supplementary Table S1**. Genotype information for PDA-XX samples has been previously published and is based on targeted sequencing approach of organoids derived from those tissues (32). The genotype for all VR-Px samples was assessed in the same way. All tissues analyzed in this study were treatment naïve.

### Mouse models

Male and female C57BL/6J (strain number 632, Charles River, RRID:IMSR_JAX:000664), athymic nude nu/nu (strain number 490, Charles River, RRID:IMSR_CRL:490) and NSG mice (strain number 614, Charles River, RRID:IMSR_JAX:005557) were purchased from the Charles River Laboratory (7-9-week-old at arrival). All animals are housed in accordance with the guidelines of the UK Home Office “Code of Practice for the Housing and Care of Experimental Animals”. They are kept behind strict barriered housing, which has maintained animals at a well-defined microbiological health status. This accommodation precludes access by wildlife, including rodent and insect vectors, and is free of infestation with ectoparasites. All animals are health screened every 3 months according to the FELASA guidelines (FELASA 2002). All animals are fed expanded rodent diet (Labdiet) and filtered water ad libitum. Environmental enrichment includes nesting material, structures for three-dimensional use of the cage and an area to retreat, and provision of chew blocks. All animal procedures and studies were reviewed by the CRUK-CI AWERB, approved by the Home Office and conducted under project license number PP4778090 in accordance with relevant institutional and national guidelines and regulations.

### Orthotopic transplantation models of PDAC

Pancreas injections were conducted as previously described (5,11). Briefly, single cells (10,000 murine cells/mouse or 30,000-700,000 human cells/mouse (700,000 cells were injected for NSG cohorts 5 and 6) prepared from PDAC organoid cultures were resuspended as a 35 μL suspension of 50% Matrigel in PBS and injected into the pancreas of 8-10-week-old mice. Pancreatic tumors were imaged using the Vevo 2100 Ultrasound at two different orientations with respect to the transducer. Tumor volumes were measured at two or three angles, whenever possible, using the Vevo LAB software program (version 5.7.0). Tumor volume analyses were performed blindly prior to plotting the data for visualization. Only mice with successful (i.e., non-leaked) orthotopic injections were included. For each experiment, SMAD4 WT and SMAD4-deficient/KO cohorts were transplanted on the same day and imaged by ultrasound on the same day. NSG mice were used for transplantation of human PDAC organoids, nu/nu mice were used for transplantation of KvPC (i.e. from the *Kras^FRT-LSL-^*^G12V*-FRT/+*^; *Trp53^LSL-^*^R172H^; *Pdx1-Cre*; *Rosa26-FlpO^ERT2^* GEMM) organoids since this GEMM had a mixed, not pure C57BL/6J, background (39). Nu/nu mice were also used for transplantation of KPC PDAC organoids that were generated by stable expression of CAS9, rather than CAS9 electroporation (see below for details about the two CRISPR technologies used to generate *Smad4* WT and *Smad4* KO organoid lines), and to enable comparison with KvPC PDAC models. C57BL/6J mice were used for all other transplant models.

### *In vivo* AZD1480 treatment study

The drug was prepared daily as a suspension in 0.1% Tween80, 0.5% hydroxyl propyl methyl cellulose in sterile water, and sonicated before administration. Palpable pancreatic tumors in C57BL/6J mice were imaged prior to enrolment (day -1) and at endpoint (day 14) using the Vevo 2100 Ultrasound at two-three different orientations, whenever possible, with respect to the transducer. Mice with tumor diameters of 5 to 8 mm were randomized and enrolled 1 day after scanning (i.e. day 0). Mice were administered vehicle or 50-60 mg/kg of AZD1480 (S2162, CAS 935666-88-9; Selleck or HY-10193, CAS 935666-88-9; MedChem Express) for 14 days, once a day (in the AM) via oral gavage. Tumor volumes were measured at two or three angles, whenever possible, using the Vevo LAB software program (version 5.7.0), and growth rate was measured by dividing the volume at day 14 for the volume at day -1. Tumor volume analyses were performed blindly prior to plotting the data for visualization.

### Cell lines and cell culture

Murine PSCs (SV40-immortalised, C57BL/6J background), apart from PSC21, and murine PDAC KPC organoid lines (C57BL/6J background) were previously described (4,34). The PSC21 line was generated and immortalized as previously described (4,11). Briefly, to establish PSC21, we utilized two and a half pancreata, and a density gradient centrifugation method with Histodenz (D2158; Sigma-Aldrich) and Gey’s Balanced Salt Solution (G9779; Sigma-Aldrich). Murine KvPC (i.e. FPC organoids from the *Kras^FRT-LSL-^*^G12V*-FRT/+*^; *Trp53^LSL-^*^R172H^; *Pdx1-Cre*; *Rosa26-FlpO^ERT2^* GEMM, not pure C57BL/6J background) organoids were kindly obtained via Material Transfer Agreement (MTA) from Professor Tuveson (Cold Spring Harbor Laboratory, CSHL) and have been previously published (39). They were then cultured for three passages in complete organoid media with 10 μM Nutlin-3a (SML0580; Sigma-Aldrich) to enrich for organoids that undergone p53 loss of heterozygosity (LOH) and generate T-LOH organoid lines, as previously done for KPC organoids (34). By inhibiting the interaction between p53 and the E3 ubiquitin protein ligase MDM2, Nutlin-3a leads to WT p53 activation, consequential depletion of cells that retain the WT *Trp53* allele and upregulation of mutant p53 levels (34,55,56). Mouse PSCs were cultured in DMEM (41966029; Gibco) containing 5% FBS. All cells were cultured for no more than 30 passages, whenever possible, at 37°C with 5% CO2. Cell line authentication was previously performed at the CRUK-CI for murine PSC4 and PSC5. Mycoplasma testing for murine PSCs was performed prior to freezing. Human organoids were kindly obtained via MTA from Professor Tuveson (CSHL) and have been previously published (33). All tissue donations to generate these organoids had been reviewed and approved by the Institutional Review Board of CSHL and all clinical institutions. Written informed consent was obtained prior to acquisition of tissue from all patients. The studies were conducted in accordance to recognized ethical guidelines (Declaration of Helsinki). Clinical information of the human PDAC organoids used is provided in **Supplementary Table S2**.

### *In vitro* cell treatments

PSCs were treated in Matrigel in 5% FBS DMEM with PDAC organoid conditioned media (CM) for as long as specified in the figure legends. KPC and KvPC PDAC organoids were treated with 8 μM AZD1480 for 48 hours prior to Western blot analysis of p-STAT3 and STAT3 levels or for 144 hours for proliferation assays.

### PCR-based genotyping of *Trp53* 1loxP

PCR-based genotyping of *Trp53* 1loxP (Figure S5A) was previously described (34). Briefly, organoids were harvested and centrifuged at 1,000 rpm for 5 min at 4°C. Genomic DNA was extracted from organoids with DNEasy Blood & Tissue Kit (69504; Qiagen). *Trp53* 1loxP genotyping PCR reaction was performed in a 20 μL reaction using AmpliTaq Gold 360 master mix (4398881; Thermo Fisher Scientific), 0.5 μM each primer (p53loxF AGCCTGCCTAGCTTCCTCAGG and p53loxR CTTGGAGACATAGCCACACTG) and 100 ng of template DNA. The PCR cycling conditions were 95°C for 10 min, followed by 40 cycles at 95°C for 30 s, 56°C for 30 s, and 72°C for 30 s, then 72°C for 7 min (BioRad T100 Thermocycler). PCR products were separated on a 2% agarose gel in TAE buffer. Gel imaging was performed with a Syngene U:Genius 3.

### *Smad4* CRISPR/Cas9 knockout

We utilized two different strategies to generate *Smad4* KO organoids.

To knock out *Smad4* in organoids for KPC T6-LOH and T69A KO 1 and KO 2 clones/pools, lenti-Cas9-Blast plasmids (52962; Addgene) for stable expression of CAS9 protein were used as previously described (5). Briefly, organoids were prepared as single cells and infected and selected using 2 μg/mL blasticidin (A11139-03; Thermo Fisher Scientific). Single guide RNAs (sgRNA) were designed using Benchling (RRID:SCR_013955) and cloned into the LRGN (LentisgRNA-EFS-GFP-neo) plasmid. Organoids were plated as single clones in the presence of geneticin (10131035; Thermo Fisher Scientific). Knockout was confirmed by western blot analysis. sgRNAs against the *Rosa26* locus were included to generate control (i.e. WT) lines.

To knock out *Smad4* in organoids for KPC T6-LOH and T69A 263 and 264 clones/pools (i.e. KO 3 and KO 4) and KvPC T93-LOH and T95-LOH 263 and 264 clones/pools (i.e. KO 3 and KO 4), CRISPR guides (phosphorothionate-modified sgRNA, 263: ATCAGGCCACCTCCACAGAC, 264: AGACGGGCATAGATCACATG) were designed against exon 3 of the murine *Smad4* gene (ENSMUST00000025393.14). A guide which targets the *Rosa26* locus was also included to generate control (i.e. WT) lines (265: GAAGATGGGCGGGAGTCTTC). Mouse PDAC organoids were dissociated into single cells and 100,000 cells were electroporated using an Amaxa 4D Nucleofector unit (Lonza) with 4 μg TrueCut spCas9 protein V2 (A36498; Invitrogen) and 80 pmol guide RNA (Synthego), using program CM-137 and P3 nucleofector solution (V4XP-3032; Lonza). A cell pellet was taken 3- and10-days post electroporation, and genomic DNA was extracted using the DNeasy blood and tissue kit (69506; Qiagen). Exon 3 of *Smad4* was amplified by PCR using the Q5 High Fidelity DNA polymerase (M0491S; NEB). *Smad4* primers used were FWD: TTCCCTTCAGCAGAAGCTGG, and REV: TGCTTCCCATACTGTTTGCA. Amplicons were subjected to Sanger sequencing and analyzed using Synthego ICE web tool to calculate the percent editing in a pool. Organoids were plated as single clones and knockout was confirmed by western blot analysis.

### Western blot analyses

PSCs and organoids were harvested in Cell Recovery Solution (354253; Corning) supplemented with complete, mini protease inhibitors (11836170001; Roche) and a phosphatase inhibitor cocktail (4906837001; Roche) and incubated for 30 min at 4°C. Cells were pelleted at 1500 rcf for 5 min and lysed in 0.1% Triton X-100, 15 mmol/L NaCl, 0.5 mmol/L EDTA, 5 mmol/L Tris, pH 7.5, supplemented with complete, mini protease inhibitors (11836170001; Roche) and a phosphatase inhibitor cocktail (4906837001; Roche). Cells were incubated on ice for 30 min briefly vortexed and pelleted at 13,200 rpm for 10 min at 4°C. Concentration of protein collected in the supernatant was determined using DC protein assay (5000113-5; Bio-Rad). Standard procedures were used for western blotting. Primary antibodies used were ACTIN (8456; Cell Signaling Technology; RRID:AB_10998774), HSP90 (07-2174;Millipore; RRID:AB_10807022), SMAD4 (sc-7966; Santa Cruz; RRID:AB_627905), p-STAT3 (9145; Cell Signaling Technology; RRID:AB_2491009), STAT3 (9139; Cell Signaling Technology; RRID:AB_331757), p53 (P53-CM5P-L; Leica; RRID:AB_2744683) and GFP (ab6673; Abcam; RRID:AB_305643). Proteins were detected using appropriate HRP-conjugated secondary antibodies (Jackson ImmunoResearch Laboratories). All western blots are representative examples and have been repeated for at least two biological replicates.

### Proliferation assays

For proliferation assays of PDAC organoids, 5,000 single cell organoids were plated in 50 μL of 100% Matrigel on 24-well plates (Corning/Nunc) and cultured in 500 μL of reduced media (5% FBS DMEM-029) or complete mouse organoid media (35). Organoid proliferation was followed for 96-144 hours with an Incucyte organoid module (Sartorius) with measurement of the organoid area per well every 3 hours (with 4 technical replicates per measurement). Data were normalized to the first measurement (at 3 hours post-plating on day 0). For proliferation assays of PDAC organoids with 8 μM of the JAKi AZD1480, 2,000-3,000 single cell organoids were plated in 25 μL of 100% Matrigel on 48-well plates (Corning/Nunc) and cultured in 250 μL of reduced media (5% FBS DMEM-029). Organoid proliferation was followed as before, data were normalized to the first measurement (i.e. average of 6 technical replicates, at 3 hours post-plating on day 0) of DMSO treated controls and used to calculate cell viability in comparison to DMSO treated controls.

### Immunohistochemical and histological analyses

Standard procedures were used for IHC. Primary antibodies for IHC were αSMA (ab5694; Abcam; RRID:AB_2223021), SMAD4 (sc-7966; Santa Cruz; RRID:AB_627905) and p-STAT3 (9145; Cell Signaling Technology; RRID:AB_2491009). Hematoxylin (H-3404-100, Vector Lab) was used as nuclear counterstain. Masson’s trichrome stains were performed according to standard protocols by the Histology core at the CRUK-CI. Stained sections were scanned with Aperio ScanScope CS and analyzed using ImageScope software (RRID:SCR_014311) Positive Pixel Count algorithms or a Nuclear v9 algorithm. Images of tissue slides were obtained with an Axio Vert.A1 (ZEISS) apart for human PDAC tumors, which were snapshotted from the ImageScope software. The percentage of collagen area was then determined by calculating the percentage of blue pixels relative to the entire stained area. To quantify αSMA stain, the percentage of positive pixels was calculated relative to the entire section. To quantify p-STAT3 stain, the percentage of positive nuclei was calculated relative to the total number of nuclei. For Masson’s trichrome and αSMA quantification of human PDAC tissues, only the PDAC area was included for analysis, following annotation from a pathologist. Human PDAC organoids hT1 and hT108 did not generate tumors following transplantation, and could not be analyzed. Quantifications were performed blindly prior to plotting the data for visualization.

### Flow cytometry analyses

Tumors were processed as previously described (5). Cells were blocked for 15 min on ice with CD16/CD32 Pure 2.4G2 (553142, BD Bioscience; RRID:AB_394657). For flow cytometric analysis of endothelial cells, immune cells, epithelial cells and iCAFs, myCAFs, CD90^+^ CAFs, CD90^-^ CAFs and apCAFs, cells were stained for 30 min on ice with anti-mouse CD31-PE/Cy7 (102418; BioLegend; RRID:AB_830757), CD45-PerCP/Cy5.5 (103132; BioLegend; RRID:AB_893344), CD326 (EpCAM)-AlexaFluor 488 (118210; BioLegend; RRID:AB_1134099), PDPN-APC/Cy7 (127418; BioLegend; RRID:AB_2629804), MHCII-BV785 (107645; BioLegend; RRID:AB_2565977), Ly6C-APC (128015; BioLegend; RRID:AB_1732087) and CD90-PE (ab24904; Abcam; RRID:AB_448474).

For flow cytometric analysis of CD56^+^ and CD49E^+^ CAFs of KPC cohorts 1 and 2 in nu/nu mice, cells were stained for 30 min on ice with anti-mouse CD56-APC (FAB7820A; Bio-Techne), CD49e-PE (557447; BD Biosciences; RRID:AB_396710), CD326 (EpCAM)-AlexaFluor 488 (118210; BioLegend; RRID:AB_1134099), PDPN-APC/Cy7 (127418; BioLegend; RRID:AB_2629804), CD45-BV785 (103149; BioLegend; RRID:AB_2564590), CD26 PerCP/Cy5.5 (45-0261-82; Thermo Fisher Scientific;, RRID:AB_1548738 – data not shown) and CD34 PE/Cy7 (25-0349-41; Thermo Fisher Scientific; RRID:AB_1963577 – data not shown). For flow cytometric analysis of CD105^+^ CAFs of KPC cohorts 1 and 2 in nu/nu mice, cells were stained for 30 min on ice with anti-mouse CD105-PE/Cy7 (120409; BioLegend; RRID:AB_1027702), CD45-PerCP/Cy5.5 (103132; BioLegend; RRID:AB_893344), CD326 (EpCAM)-AlexaFluor 488 (118210; BioLegend; RRID:AB_1134099), PDPN-APC/Cy7 (127418; BioLegend; RRID:AB_2629804), Ly6C BV785 (128041;BioLegend; RRID:AB_2565852 – data not shown) and CD140a PE (135906; BioLegend; RRID:AB_1953269 – data not shown).

For flow cytometric analysis of CD56^+^, CD49E^+^ and CD105^+^ CAFs of KPC cohort 3 in nu/nu mice and all other KPC and KvPC cohorts, cells were stained for 30 min on ice with anti-mouse CD56-APC (FAB7820A; Bio-Techne), CD49e-PE (557447; BD Biosciences; RRID:AB_396710), CD105-PE/Cy7 (120409; BioLegend; RRID:AB_1027702), CD326 (EpCAM)-AlexaFluor 488 (118210; BioLegend; RRID:AB_1134099), PDPN-APC/Cy7 (127418; BioLegend; RRID:AB_2629804), CD45-BV785 (103149; BioLegend; RRID:AB_2564590) and CD26 PerCP/Cy5.5 (45-0261-82; Thermo Fisher Scientific; RRID:AB_1548738 – data not shown).

For flow-cytometric analysis of macrophages and neutrophils of all KPC and KvPC cohorts, cells were stained for 30 min on ice with anti-mouse CD45-PerCP/Cy5.5 (103132; BioLegend; RRID:AB_893344), CD11b-PE/Cy7 (101215; BioLegend; RRID:AB_312798), Ly6c-Alexa488 (128021; BioLegend; RRID:AB_10640820 – data not shown), F4/80-BV785 (123141; BioLegend; RRID:AB_2563667), MHCII-APC/Cy7 (107627; BioLegend; RRID:AB_1659252 – data not shown), CD11c-APC (117309; BioLegend; RRID:AB_313779 – data not shown), Gr1-PE (108407; BioLegend; RRID:AB_313372).

Cells were resuspended in PBS with DAPI and analyzed on a BD FACSymphony cell analyzer.

Flow analyses were performed blindly using FlowJo 10.8.2 (RRID:SCR_008520) prior to plotting the data for visualization.

### Cell sorting of PDAC organoid/PSC co-cultures for RNA-sequencing

Sorting of PDAC organoid/PSC co-cultures was performed following 3.5 days culture in 5% FBS DMEM. Following single cell digestion of co-cultures, cells were stained for 30 minutes with anti-mouse CD326 (EpCAM)-PE (118205; BioLegend; RRID:AB_1134176) and PDPN-AlexaFluor 488 (156208; BioLegend; RRID:AB_2814080). Cells were resuspended in PBS with DAPI and sorted with a BD FACSMelody cell sorter.

### RNA-sequencing analyses of PDAC organoids and PSCs flow-sorted from co-cultures

RNA-seq data of KPC monocultures and co-cultures are available at the Gene Expression Omnibus (GEO) under the accession number GSE263080. RNA-seq data of KvPC monocultures and co-cultures are available at the GEO under the accession number GSE263081. Samples were collected in 1 mL of TRIzol Reagent (15596018; Invitrogen). RNA was extracted using the PureLink RNA mini kit (12183018A; Invitrogen). RNA concentration was measured using a Qubit and RNA quality was assessed on a TapeStation 4200 (Agilent) using the Agilent RNA ScreenTape kit. mRNA library preparations were performed using 55 μL of 10 ng/mL per sample (RNA integrity number > 8, except for one sample with RIN 7.2). Illumina libraries were then sequenced on 1 lane of SP flowcell on NovaSeq6000. FASTQ files were aligned, and the expression levels of each transcript were quantified using Salmon (v1.4.0) (57) with the annotation from ENSEMBL (GRCh38 release 98) with recommended settings. Transcript-level expression was loaded and summarized to the gene level by using tximport (58). Differential gene expression was performed using DESeq2 (v2) (59) by applying lfcshrink function (60). The principal components for variance-stabilized data were estimated using plotPCA function, available in DESeq, and ggplot2 (https://ggplot2.tidyverse.org). Genes with adjusted p-value < 0.05 were selected as significantly differentiated between conditions. Following differential gene expression, genes were pre-ranked based on the negative logarithmic p-value and the sign of the log2 fold change. GSEA was performed using clusterprofiler against the Hallmark, Reactome, and C2 canonical pathway collection (C2.cp.v5.1) downloaded from the Molecular Signatures Database (MSigDB) (61). GSEA plots were plotted using enrichplot R package (https://bioconductor.org/packages/enrichplot).

***NicheNet:*** NichenetR (38) was used to infer the ligands activity and their regulation potential of a sender group by considering the expression of downstream genes in the receiver group (i.e. target genes). Ligands were ranked based on their area under the precision-recall curve (AUPR) to prioritize ligands that regulate the target genes in the receiver population. Nichenet was applied on bulk RNA-seq co-culture data to infer the interaction between PDAC organoids and PSCs in KPC *Smad4* WT and KPC *Smad4* KO conditions. For Nichenet, “target-genes” were defined by using significant DEG (p.adj < 0.05, log2FC > 1) in each condition of the receiver group, and the “ligands” were defined the same way in the sender group. All expressed genes in the dataset were considered as “background expressed genes”.

### Single-cell RNA-sequencing analyses of PDAC tumors

scRNA-seq data of KPC tumors are available at the GEO under the accession number GSE262879. scRNA-seq data of KvPC tumors are available at the GEO under the accession number GSE262878. Cell Ranger (10x Genomics) workflow (62) was used to align FASTQ files to GRCh38 (mm10) mouse transcriptome reference to generate the raw counts of gene expression quantities. The reference genome was modified by adding *Gfp* to it. From the raw counts of each sample, the SOLO tool that is implemented in scvi-tools was used to estimate doublets to be removed (63). The samples were integrated, and batch effect was removed using scvi-tools (64). Scanpy workflow (65) was used for dimension reductions, clustering and defining markers of each cluster. Cells that have the percentage of mitochondrial genes more than 5% were filtered out. Unique molecular identifiers (UMI) were normalized to 10,000 counts and Leiden graph clustering were used for unsupervised clustering to identify the cell populations with similar transcriptomic profile. For dimension reduction we built the top 30 principal components (PC) and nearest neighbors’ graph (k = 10) on 2,000 highly variable genes. Uniform manifold approximation and projection (UMAP) were used to visualize the datasets in 2-dimentional space. Markers of each cluster were defined using “rank_genes_groups” function from Scanpy.

***Copy Number Varation (CNV) analysis:*** A python implementation of inferCNV of the Trinity CTAT Project (https://github.com/broadinstitute/inferCNV) was used to estimate the copy number status in each cell type. We used the fibroblast cells as a reference key and a 250-genes window size.

***Pseudobulk:*** For DE analysis we pooled all cells within a specific cell type by summing the gene expression of each gene to create a pseudo-bulk expression profile of each sample. DESeq2 uses the pseudo-bulk data to detect the differences between two sample groups. To generate the pseudo-bulk profile and perform DEA, we used the python implementation of decoupler R package (66) with the default options. GSEA was performed as with bulk RNA-seq data.

***CellChat:*** Cellchat R package (37) was used with the recommended setting to infer and visualize the cell-cell communication between specific cell types in scRNA-seq data.

### Statistical analysis

GraphPad Prism software (RRID:SCR_002798), customized R and python scripts were used for graphical representation of data. Statistical analysis was performed using non-parametric Mann-Whitney test or chi-square test. All statistical details of experiments are specified in the figure legends and/or panel figures, including the number of technical and biological replicates, and how significance was defined.

## Data availability

For RNA-seq and scRNA-seq datasets, the data generated in this study are publicly available in Gene Expression Omnibus (GEO) at GSE263080, GSE263081, GSE262879 and GSE262878. For ultrasound-based tumor volume, western blot, organoid proliferation assay, flow cytometry and immunochemistry data, the raw data are available on reasonable request from the corresponding author. No publicly available data was reused. This paper does not report original code.

## Supporting information

Combined Supplementary Figures and Legends

## ACKNOWLEDGEMENTS

The authors would like to thank the BRU, Genomics, Bioinformatics, Flow Cytometry, Pre-genome editing and Histology core facilities at the Cancer Research UK Cambridge Institute (CRUK-CI). This work was mainly supported by a UKRI Future Leaders Fellowship of which GB is recipient and that also supports W.L. This work was also supported by a Cancer Research UK institutional grant (A27463), which also supported J.S.M., S.A. and S.P.T. Also, G.B. is recipient of a Pancreatic Cancer Research Foundation grant and a US Department of Defense PCARP grant, which supported G.M. and J.S.M., a NCI-CRUK Cancer Grand Challenge grant, which supported M.J., W.K.L. and S.H., and a Pancreatic Cancer UK Future Leaders Academy grant, which supported P.S.W.C. E.G.L. was supported by a MRC Doctoral Training Grant. J.A.H. was supported by a Harding Distinguished Postgraduate Programme PhD studentship (Cambridge Trust). The authors would like to thank Professor David Tuveson for sharing the human PDAC organoids employed in this study, as well as the FPC organoids.

## AUTHOR CONTRIBUTIONS

E.G.L. designed and conducted the experiments, and wrote the paper. S.P.T, J.S.M., M.J., J.A.H, W.L., W.K.L, S.H., P.S.W.C., M.Z., P.M.J. and L.V. conducted the experiments. R.B. performed histopathology analysis of murine and human tissues. V.C. and A.S. shared human PDAC tissues. M.V. coordinated the *in vivo* mouse work. G.B conceptualized and supervised the study, designed and conducted the experiments, and wrote the paper.

