## Supplementary material for "SMAD4 and KRAS status shape malignant-stromal crosstalk in pancreatic cancer": Combined Supplementary Figures and Legends

### SUPPLEMENTARY FIGURE LEGENDS

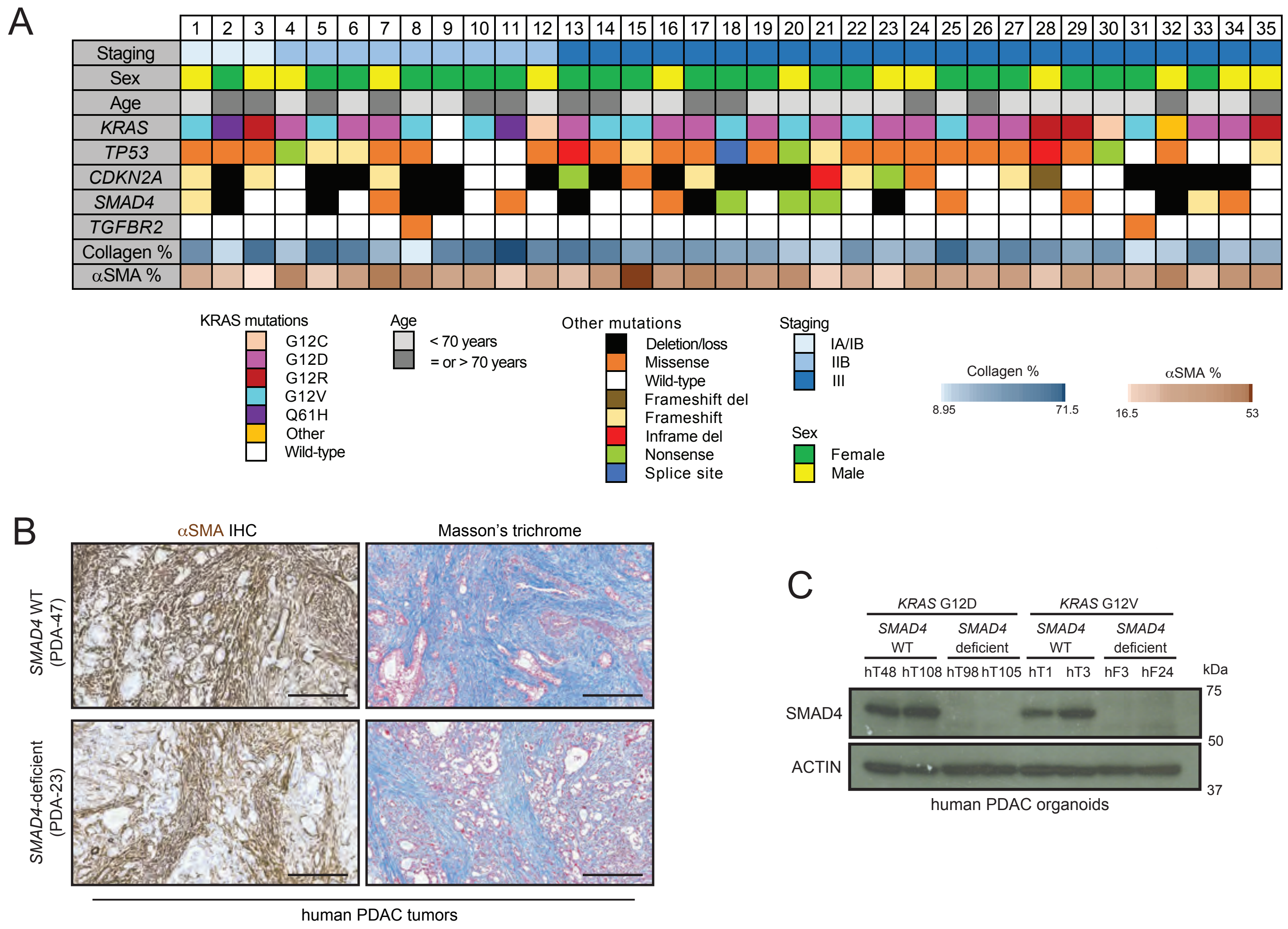


D

| Mouse cohorts | Organoid lines | KRAS status | TP53 status | SMAD4 status | CDKN2A status |
|---|---|---|---|---|---|
| 1-2 ♀/♂ NSG | hT1 | G12V | R213* | WT | Deep loss |
| | hF24 | G12V | F113V | Loss | Deep loss |
| 3-4 ♀/♂ NSG | hT108 | G12D | P98fs | WT | D108fs |
| | hT98 | G12D | M237I | Deep loss | R80* |
| 5 ♂NSG | hT3 | G12V | Loss | WT | Deep loss |
| | hF3 | G12V | R273L | Deep loss | R58* |
| 6 ♂NSG | hT48 | G12D | P151T | WT | Loss |
| | hT105 | G12D | P142fs | Deep loss | Deep loss |

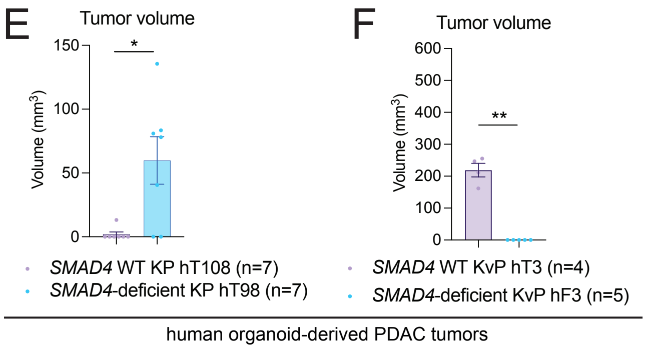


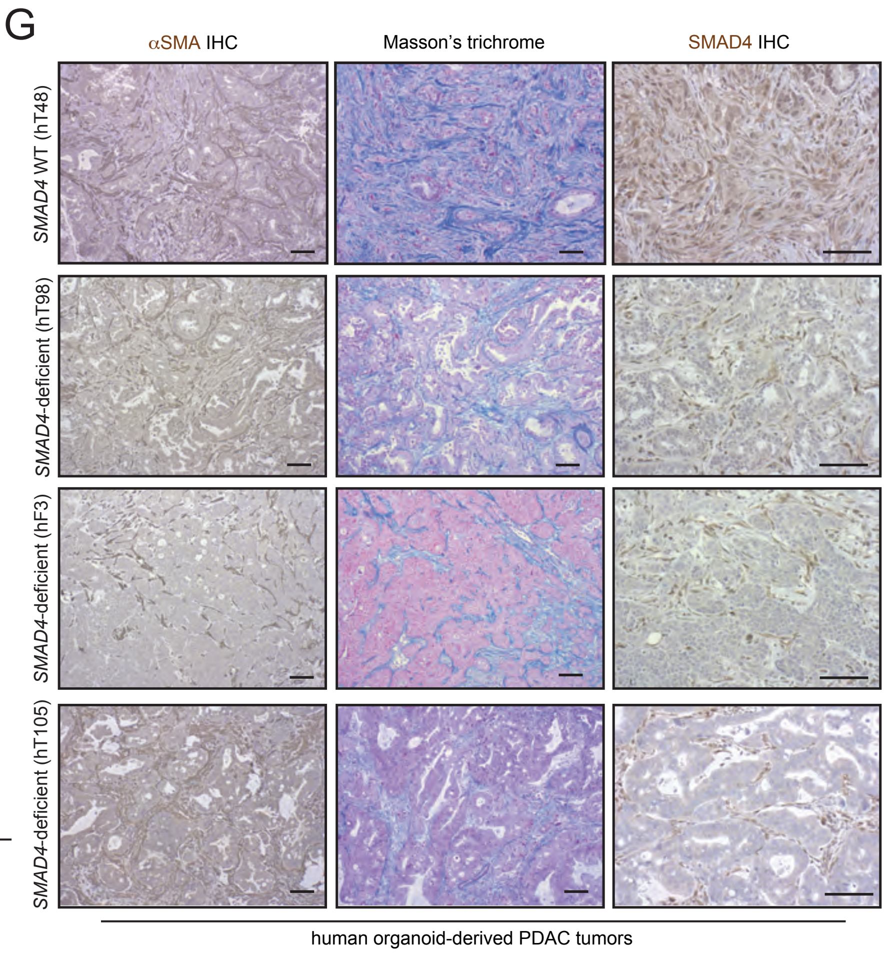


Figure S1

**Figure S1. *SMAD4*-deficient human organoid-derived PDAC tumors have less fibrosis than *SMAD4* WT tumors. (A)** Summary of clinical information, genetic alterations, collagen abundance and alpha smooth muscle actin (αSMA) abundance of a panel of human PDAC tumors (n=35). Masson's trichrome (for collagen) and αSMA quantification were calculated as % positive area / total PDAC area following annotation of the tissues by a pathologist. **(B)** Representative Masson's trichrome and αSMA stains in *SMAD4* WT or *SMAD4*-deficient human PDAC tumors. Scale bars, 300 μm. **(C)** Western blot analysis of SMAD4 in *SMAD4* WT or *SMAD4*-deficient human PDAC organoids with $KRAS^{G12D}$ (KP) or $KRAS^{G12V}$ (KvP) mutation. ACTIN, loading control. **(D)** Table summarizing the experimental cohorts of orthotopic transplantation models of KvP and KP human PDAC organoids in NOD scid gamma (NSG) mice. **(E-F)** Tumor volumes as measured by ultrasound-based imaging of tumors derived from the transplantation of *SMAD4* WT or *SMAD4*-deficient KP **(E)** or KvP **(F)** human PDAC organoids with $KRAS^{G12D}$ or $KRAS^{G12V}$ mutation, respectively. Results show mean ± SEM (n=4-7 mice/cohort). *, $P < 0.05$; **, $P < 0.01$, Mann-Whitney test. **(G)** Representative Masson's trichrome, SMAD4 and αSMA stains in PDAC tumors derived from the orthotopic transplantation of *SMAD4* WT or *SMAD4*-deficient KP or KVP human PDAC organoids. Scale bars, 50 μm.

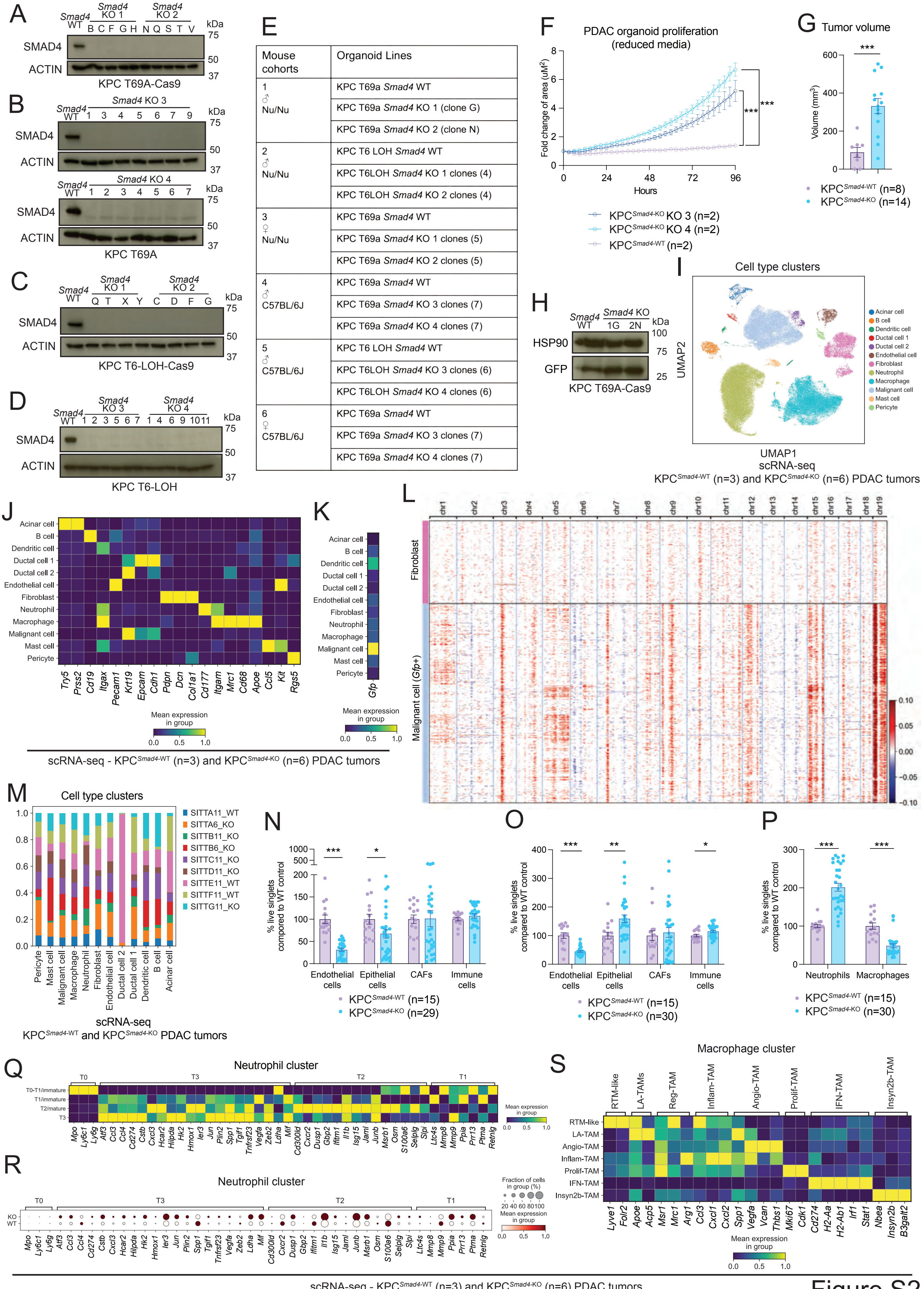
A
Smad4 WT
Smad4 KO 1
Smad4 KO 2
B C F G H N Q S T V
kDa
75
50
37
SMAD4
ACTIN
KPC T69A-Cas9
B
Smad4 KO 3
Smad4 KO 4
KPC T69A
C
KPC T6-LOH-Cas9
D
KPC T6-LOH
E
F
PDAC organoid proliferation (reduced media)
Fold change of area (uM²)
Hours
KPC Smad4-KO KO 3 (n=2)
KPC Smad4-KO KO 4 (n=2)
KPC Smad4-WT (n=2)
G
Tumor volume
Volume (mm³)
KPC Smad4-WT (n=8)
KPC Smad4-KO (n=14)
H
HSP90
GFP
Smad4 WT
Smad4 KO
1G 2N
KPC T69A-Cas9
I
Cell type clusters
UMAP1
UMAP2
scRNA-seq
KPC Smad4-WT (n=3) and KPC Smad4-KO (n=6) PDAC tumors
Acinar cell
B cell
Dendritic cell
Ductal cell 1
Ductal cell 2
Endothelial cell
Fibroblast
Neutrophil
Macrophage
Malignant cell
Mast cell
Pericyte
J
Try5 Prss2 Cd19 Itgax Pecam1 Krt19 Epcam Cdh1 Pdpn Dcn Col1a1 Cd177 Itgam Mrc1 Cd68 Apoe Ccl5 Kit Rgs5
Mean expression in group
K
Gfp
scRNA-seq - KPC Smad4-WT (n=3) and KPC Smad4-KO (n=6) PDAC tumors
L
Fibroblast
Malignant cell (Gfp+)
chr1 chr2 chr3 chr4 chr5 chr6 chr7 chr8 chr9 chr10 chr11 chr12 chr13 chr14 chr15 chr16 chr17 chr18 chr19
M
Cell type clusters
SITTA11_WT
SITTA6_KO
SITTB11_KO
SITTB6_KO
SITTC11_KO
SITTD11_KO
SITTE11_WT
SITTF11_WT
SITTG11_KO
scRNA-seq
KPC Smad4-WT and KPC Smad4-KO PDAC tumors
N
% live singlets compored to WT control
Endothelial cells
Epithelial cells
CAFs
Immune cells
KPC Smad4-WT (n=15)
KPC Smad4-KO (n=29)
O
% live singlets compared to WT control
KPC Smad4-WT (n=15)
KPC Smad4-KO (n=30)
P
Neutrophils
Macrophages
Q
Neutrophil cluster
T0 T3 T2 T1
T0-T1/immature
T1/immature
T2/mature
T3
R
Neutrophil cluster
Fraction of cells in group (%)
KO
WT
S
Macrophage cluster
RTM-like
LA-TAMs
Reg-TAM
Inflam-TAM
Angio-TAM
Prolif-TAM
IFN-TAM
Insyn2b-TAM
Mean expression in group

Figure S2

**Figure S2. *Smad4* loss impacts the immune TME in KPC PDAC. (A-D)** Validation of *Smad4* KO KPC PDAC organoids (n=2 parental lines) by western blot analysis of SMAD4 in KPC$^{Smad4\text{-}WT}$ (i.e. *Rosa26* KO) pool or KPC$^{Smad4\text{-}KO}$ clones (from 3 different guides with 2 different CRISPR/Cas9-based methods) cultured in complete organoid media. ACTIN, loading controls. **(E)** Table summarizing the experimental cohorts of orthotopic transplantation models of KPC$^{Smad4\text{-}WT}$ and KPC$^{Smad4\text{-}KO}$ organoids in nu/nu or C57BL/6J mice. **(F)** Proliferation curves of KPC$^{Smad4\text{-}WT}$ and KPC$^{Smad4\text{-}KO}$ organoids cultured for 96 hours in Matrigel in reduced media. Data were normalized to the first measurement (at 3 h post-plating on day 0). Results show mean $\pm$ SEM of n=2 biological replicates (with n=4 technical replicates each). ***, $P < 0.001$, Mann-Whitney test calculated for the last time point. **(G)** Tumor volumes as measured by ultrasound-based imaging of tumors derived from the transplantation of KPC$^{Smad4\text{-}WT}$ or KPC$^{Smad4\text{-}KO}$ organoids in C57BL/6J mice. Results show mean $\pm$ SEM from 2 separate experiments, each with 1 WT group and 2 groups of KO pools from 2 different guides (n=8-14 mice/cohort, at day 32 (experiment 1) or day 21 (experiment 2) post-transplant). ***, $P < 0.001$, Mann-Whitney test. **(H)** Western blot analysis of green fluorescent protein (GFP) in KPC$^{Smad4\text{-}WT}$ pool or KPC$^{Smad4\text{-}KO}$ clones (from 2 different guides) cultured in complete organoid media. HSP90, loading control. **(I)** Uniform manifold approximation and projection (UMAP) plot shows the cell clusters from KPC$^{Smad4\text{-}WT}$ (n=3) or KPC$^{Smad4\text{-}KO}$ (n=6) tumors analyzed by single-cell RNA-sequencing (scRNA-seq). Different cell type clusters are color coded. **(J)** Heatmap of scaled expression of cell type-specific markers in each cell cluster from KPC$^{Smad4\text{-}WT}$ (n=3) or KPC$^{Smad4\text{-}KO}$ (n=6) PDAC tumors, as analyzed by scRNA-seq. Data are scaled such that the cluster with the lowest average expression = 0 and the highest = 1 for each gene. **(K)** Heatmap of scaled expression of *Gfp* expression in each cell cluster of PDAC tumors from KPC$^{Smad4\text{-}WT}$ or KPC$^{Smad4\text{-}KO}$ tumors, as analyzed by scRNA-seq. Data are scaled such that the cluster with the lowest average expression = 0 and the highest = 1 for each gene. **(L)** Heatmap showing large-scale copy number variation (CNV) profile of the fibroblast and malignant cell clusters identified by scRNA-seq. The color coding represents the CNV level based on a sliding window of 250 gene expression. Amplifications are shown in red and deletions are shown in blue. The fibroblast cluster was used as reference cell cluster. **(M)** Tumor sample contribution to different cell types in KPC$^{Smad4\text{-}WT}$ or KPC$^{Smad4\text{-}KO}$ tumors, represented as bar plots showing proportions of the different tumor samples in each cell cluster. **(N)** Flow cytometric analysis of endothelial cells ($CD31^{+}CD45^{-}$), epithelial cells ($CD45^-CD31^-EpCAM^+$), CAFs ($CD45^-CD31^-EpCAM^-PDPN^+$) and immune cells ($CD45^+$) from live singlets in $KPC^{\textit{Smad4}\text{-WT}}$ or $KPC^{\textit{Smad4}\text{-KO}}$ tumors in nu/nu mice. Results show mean ± SEM from 3 separate experiments, each with 1 WT group and 2 groups of KO pools from 2 different guides. *, $P < 0.05$, ***, $P < 0.001$, Mann-Whitney test. **(O)** Flow cytometric analysis of endothelial cells ($CD31^+CD45^-$), epithelial cells ($CD45^-CD31^-EpCAM^+$), CAFs ($CD45^-CD31^-EpCAM^-PDPN^+$) and immune cells ($CD45^+$) from live singlets in $KPC^{\textit{Smad4}\text{-WT}}$ or $KPC^{\textit{Smad4}\text{-KO}}$ tumors in C57BL/6J mice. Results show mean ± SEM from 3 separate experiments, each with 1 WT group and 2 groups of KO pools from 2 different guides. *, $P < 0.05$; **, $P < 0.01$; ***, $P < 0.001$, Mann-Whitney test. **(P)** Flow cytometric analysis of neutrophils ($CD45^+CD11b^+Gr1^+$) and macrophages ($CD45^+Gr1^-CD11b^+F4/80^+$) from live singlets in $KPC^{\textit{Smad4}\text{-WT}}$ or $KPC^{\textit{Smad4}\text{-KO}}$ tumors in C57BL/6J mice. Results show mean ± SEM from 3 separate experiments, each with 1 WT group and 2 groups of KO pools from 2 different guides. ***, $P < 0.001$, Mann-Whitney test. **(Q)** Heatmap of scaled expression of neutrophil markers in distinct neutrophil sub-clusters from $KPC^{\textit{Smad4}\text{-WT}}$ and $KPC^{\textit{Smad4}\text{-KO}}$ tumors analyzed by scRNA-seq. Data are scaled such that the cluster with the lowest average expression = 0 and the highest = 1 for each gene. **(R)** Dot plot visualization of the scaled average expression of neutrophil subset markers in neutrophils from $KPC^{\textit{Smad4}\text{-WT}}$ or $KPC^{\textit{Smad4}\text{-KO}}$ tumors analyzed by scRNA-seq. The color intensity represents the expression level and the size of the dots represents the percentage of expressing cells. **(S)** Heatmap of scaled expression of macrophage markers in distinct macrophage sub-clusters from $KPC^{\textit{Smad4}\text{-WT}}$ and $KPC^{\textit{Smad4}\text{-KO}}$ tumors analyzed by scRNA-seq. Data are scaled such that the cluster with the lowest average expression = 0 and the highest = 1 for each gene.

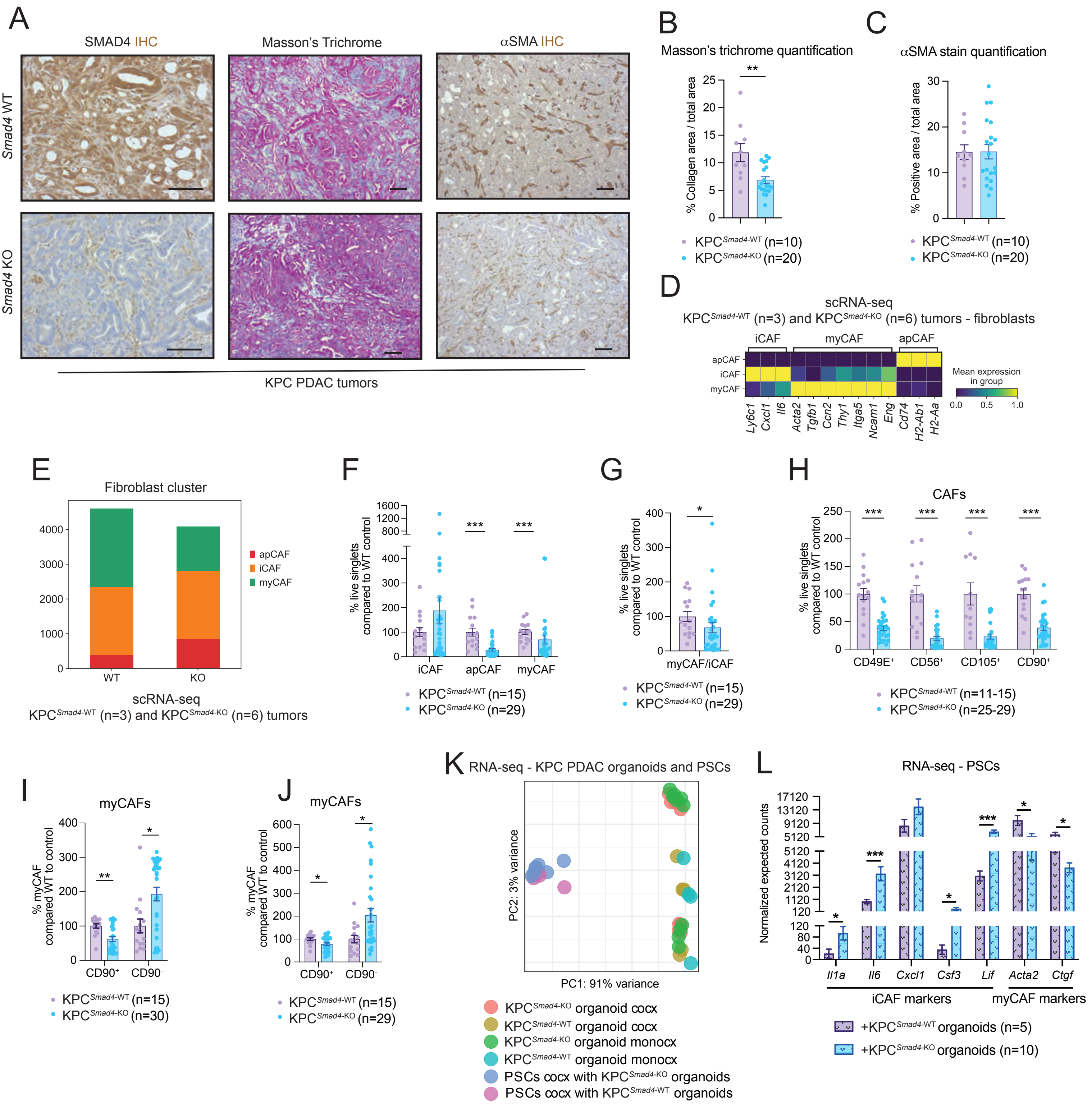
A
SMAD4 IHC
Masson's Trichrome
αSMA IHC
Smad4 WT
Smad4 KO
KPC PDAC tumors
B
Masson's trichrome quantification
% Collagen area / total area
KPC^Smad4-WT (n=10)
KPC^Smad4-KO (n=20)
C
αSMA stain quantification
% Positive area / total area
KPC^Smad4-WT (n=10)
KPC^Smad4-KO (n=20)
D
scRNA-seq
KPC^Smad4-WT (n=3) and KPC^Smad4-KO (n=6) tumors - fibroblasts
iCAF
myCAF
apCAF
Ly6c1
Cxcl1
Il6
Acta2
Tgfb1
Ccn2
Thy1
Itga5
Ncam1
Eng
Cd74
H2-Ab1
H2-Aa
Mean expression in group
E
Fibroblast cluster
apCAF
iCAF
myCAF
WT
KO
scRNA-seq
KPC^Smad4-WT (n=3) and KPC^Smad4-KO (n=6) tumors
F
% live singlets compared to WT control
iCAF
apCAF
myCAF
KPC^Smad4-WT (n=15)
KPC^Smad4-KO (n=29)
G
% live singlets compared to WT control
myCAF/iCAF
KPC^Smad4-WT (n=15)
KPC^Smad4-KO (n=29)
H
CAFs
% live singlets compared to WT control
CD49E+
CD56+
CD105+
CD90+
KPC^Smad4-WT (n=11-15)
KPC^Smad4-KO (n=25-29)
I
myCAFs
% myCAF compared WT to control
CD90+
CD90-
KPC^Smad4-WT (n=15)
KPC^Smad4-KO (n=30)
J
myCAFs
% myCAF compared WT to control
CD90+
CD90-
KPC^Smad4-WT (n=15)
KPC^Smad4-KO (n=29)
K
RNA-seq - KPC PDAC organoids and PSCs
PC2: 3% variance
PC1: 91% variance
KPC^Smad4-KO organoid cocx
KPC^Smad4-WT organoid cocx
KPC^Smad4-KO organoid monocx
KPC^Smad4-WT organoid monocx
PSCs cocx with KPC^Smad4-KO organoids
PSCs cocx with KPC^Smad4-WT organoids
L
RNA-seq - PSCs
Normalized expected counts
Il1a
Il6
Cxcl1
Csf3
Lif
Acta2
Ctgf
iCAF markers
myCAF markers
+KPC^Smad4-WT organoids (n=5)
+KPC^Smad4-KO organoids (n=10)

Figure S3

**Figure S3.** ***Smad4*** **loss drives a fibro-inflammatory stroma in KPC PDAC. (A)** Representative SMAD4, Masson's trichrome and $\alpha$SMA stains in KPC$^{Smad4\text{-WT}}$ or KPC$^{Smad4\text{-KO}}$ PDAC tumors in nu/nu mice. Scale bars, 50 $\mu$m. **(B)** Quantification of Masson's trichrome stain in KPC$^{Smad4\text{-WT}}$ or KPC$^{Smad4\text{-KO}}$ tumors in nu/nu mice. Results show mean $\pm$ SEM from 2 separate experiments, each with 1 WT group and 2 groups of KO pools from 2 different guides. **, $P < 0.01$, Mann-Whitney test. **(C)** Quantification of $\alpha$SMA stain in KPC$^{Smad4\text{-WT}}$ or KPC$^{Smad4\text{-KO}}$ tumors in nu/nu mice. Results show mean $\pm$ SEM from 2 separate experiments, each with 1 WT group and 2 groups of KO pools from 2 different guides. No statistical difference was found, as calculated by Mann-Whitney test. **(D)** Heatmap of scaled expression of CAF subtype markers in distinct CAF populations from KPC$^{Smad4\text{-WT}}$ (n=3) and KPC$^{Smad4\text{-KO}}$ (n=6) tumors analyzed by scRNA-seq. Data are scaled such that the cluster with the lowest average expression = 0 and the highest = 1 for each gene. **(E)** CAF sub-cluster abundance in fibroblasts of KPC$^{Smad4\text{-WT}}$ or KPC$^{Smad4\text{-KO}}$ tumors, as assessed by scRNA-seq. **(F)** Flow cytometric analysis of myCAFs (Ly6C$^{-}$MHCII$^{-}$), iCAFs (Ly6C$^{+}$MHCII$^{-}$) and apCAFs (Ly6C$^{-}$MHCII$^{+}$) from live singlets in KPC$^{Smad4\text{-WT}}$ or KPC$^{Smad4\text{-KO}}$ tumors in nu/nu mice. Results show mean $\pm$ SEM from 3 separate experiments, each with 1 WT group and 2 groups of KO pools from 2 different guides. ***, $P < 0.001$, Mann-Whitney test. **(G)** Flow cytometric analysis of myCAF/iCAF ratio from live singlets in KPC$^{Smad4\text{-WT}}$ or KPC$^{Smad4\text{-KO}}$ tumors in nu/nu mice. Results show mean $\pm$ SEM from 3 separate experiments, each with 1 WT group and 2 groups of KO pools from 2 different guides. *, $P < 0.05$, Mann-Whitney test. **(H)** Flow cytometric analyses of CD90$^{+}$, CD49E$^{+}$, CD56$^{+}$ and CD105$^{+}$ CAFs from live singlets in KPC$^{Smad4\text{-WT}}$ or KPC$^{Smad4\text{-KO}}$ tumors in nu/nu mice. Results show mean $\pm$ SEM from 3 separate experiments, each with 1 WT group and 2 groups of KO pools from 2 different guides. ***, $P < 0.001$, Mann-Whitney test. **(I)** Flow cytometric analysis of CD90$^{-}$ and CD90$^{+}$ myCAFs (Ly6C$^{-}$MHCII$^{-}$) from the parental gate in KPC$^{Smad4\text{-WT}}$ or KPC$^{Smad4\text{-KO}}$ tumors in C57BL/6J mice. Results show mean $\pm$ SEM from 3 separate experiments, each with 1 WT group and 2 groups of KO pools from 2 different guides. *, $P < 0.05$; **, $P < 0.01$, Mann-Whitney test. **(J)** Flow cytometric analysis of CD90$^{-}$ and CD90$^{+}$ myCAFs (Ly6C$^{-}$MHCII$^{-}$) from the parental gate in KPC$^{Smad4\text{-WT}}$ or KPC$^{Smad4\text{-KO}}$ tumors in nu/nu mice. Results show mean $\pm$ SEM from 3 separate experiments, each with 1 WT group and 2 groups of KO pools from 2 different guides. *, $P < 0.05$, Mann-Whitney test. **(K)** Principal component analysis (PCA) of PSCs and KPC PDAC organoids flow-sorted from co-cultures and monocultures analyzed by RNA-sequencing (RNA-seq). **(L)** RNA-seq expression of iCAF and myCAF markers in PSCs flow-sorted from co-cultures with KPC$^{Smad4\text{-WT}}$ or KPC$^{Smad4\text{-KO}}$ organoids. Results show mean ± SEM. *, $P < 0.05$; ***, $P < 0.001$, Mann-Whitney test.

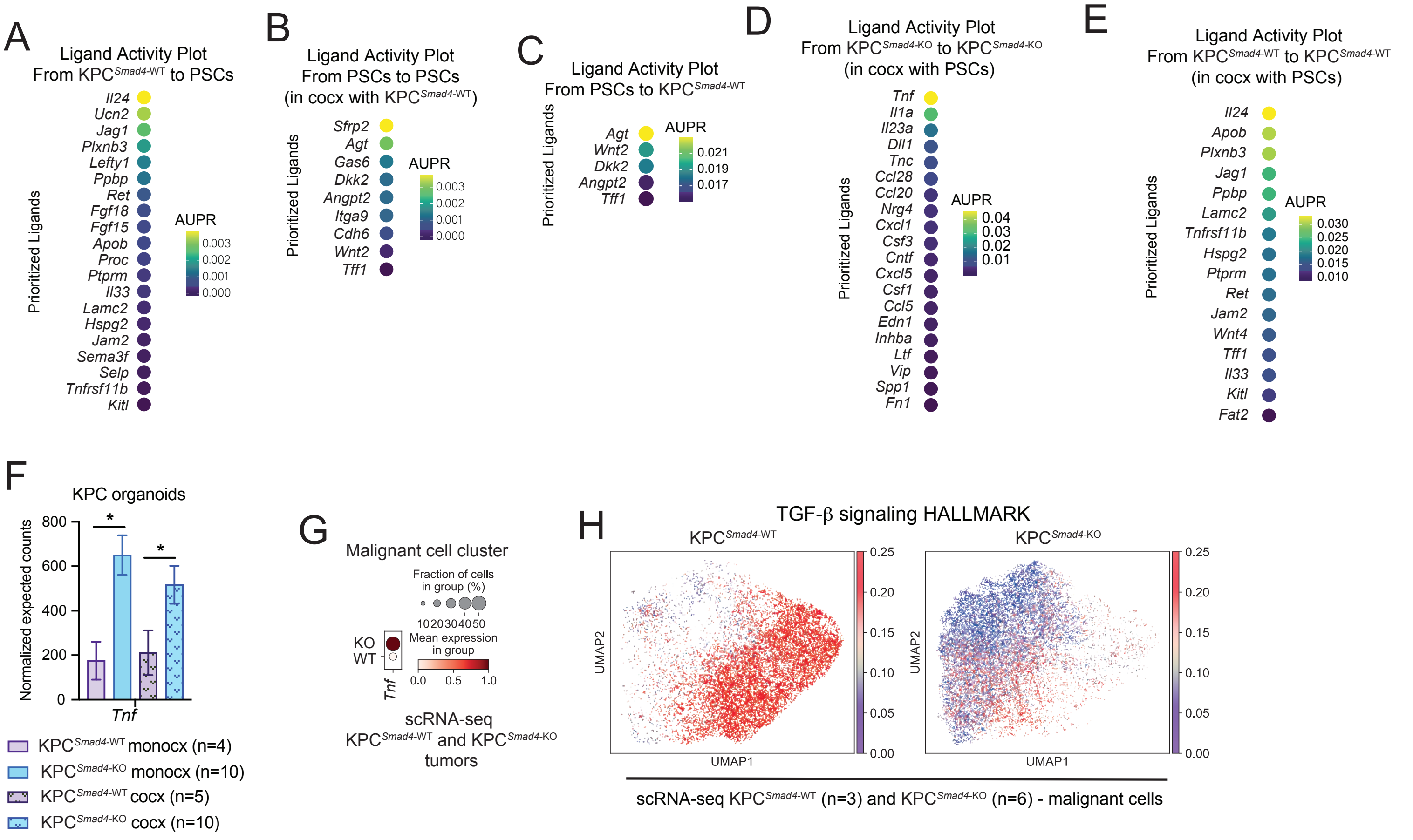
A
Ligand Activity Plot
From KPC Smad4-WT to PSCs
Prioritized Ligands
Il24
Ucn2
Jag1
Plxnb3
Lefty1
Ppbp
Ret
Fgf18
Fgf15
Apob
Proc
Ptprm
Il33
Lamc2
Hspg2
Jam2
Sema3f
Selp
Tnfrsf11b
Kitl
AUPR
0.003
0.002
0.001
0.000
B
Ligand Activity Plot
From PSCs to PSCs
(in cocx with KPC Smad4-WT)
Prioritized Ligands
Sfrp2
Agt
Gas6
Dkk2
Angpt2
Itga9
Cdh6
Wnt2
Tff1
AUPR
0.003
0.002
0.001
0.000
C
Ligand Activity Plot
From PSCs to KPC Smad4-WT
Prioritized Ligands
Agt
Wnt2
Dkk2
Angpt2
Tff1
AUPR
0.021
0.019
0.017
D
Ligand Activity Plot
From KPC Smad4-KO to KPC Smad4-KO
(in cocx with PSCs)
Prioritized Ligands
Tnf
Il1a
Il23a
Dll1
Tnc
Ccl28
Ccl20
Nrg4
Cxcl1
Csf3
Cntf
Cxcl5
Csf1
Ccl5
Edn1
Inhba
Ltf
Vip
Spp1
Fn1
AUPR
0.04
0.03
0.02
0.01
E
Ligand Activity Plot
From KPC Smad4-WT to KPC Smad4-WT
(in cocx with PSCs)
Prioritized Ligands
Il24
Apob
Plxnb3
Jag1
Ppbp
Lamc2
Tnfrsf11b
Hspg2
Ptprm
Ret
Jam2
Wnt4
Tff1
Il33
Kitl
Fat2
AUPR
0.030
0.025
0.020
0.015
0.010
F
KPC organoids
Normalized expected counts
800
600
400
200
0
*
*
Tnf
KPC Smad4-WT monocx (n=4)
KPC Smad4-KO monocx (n=10)
KPC Smad4-WT cocx (n=5)
KPC Smad4-KO cocx (n=10)
G
Malignant cell cluster
Fraction of cells in group (%)
10 20 30 40 50
Mean expression in group
0.0 0.5 1.0
KO
WT
Tnf
scRNA-seq
KPC Smad4-WT and KPC Smad4-KO tumors
H
TGF-β signaling HALLMARK
KPC Smad4-WT
KPC Smad4-KO
UMAP2
UMAP1
0.25
0.20
0.15
0.10
0.05
0.00
scRNA-seq KPC Smad4-WT (n=3) and KPC Smad4-KO (n=6) - malignant cells

Figure S4

**Figure S4. *Smad4* loss upregulates IL-1 and JAK/STAT signaling in KPC PDAC. (A)** Ligand activity plot shows the top ligands from KPC$^{Smad4\text{-WT}}$ organoids that regulate target genes in co-cultured PSCs, as assessed by NicheNet analysis of RNA-seq. AUPR, area under the precision-recall curve. **(B)** Ligand activity plot shows the top ligands from PSCs co-cultured with KPC$^{Smad4\text{-WT}}$ organoids that regulate target genes in PSCs, as assessed by NicheNet analysis of RNA-seq. **(C)** Ligand activity plot shows the top ligands from PSCs that regulate target genes in co-cultured KPC$^{Smad4\text{-WT}}$ PDAC organoids, as assessed by NicheNet analysis of RNA-seq. **(D)** Ligand activity plot showing the top ligands from KPC$^{Smad4\text{-KO}}$ organoids in co-culture with PSCs that regulate target genes in organoids, as assessed by NicheNet analysis of RNA-seq. **(E)** Ligand activity plot showing the top ligands from KPC$^{Smad4\text{-WT}}$ organoids in co-culture with PSCs that regulate target genes in organoids, as assessed by NicheNet analysis of RNA-seq. **(F)** RNA-seq expression of *Tnf* in KPC PDAC organoids flow-sorted from monocultures and co-cultures with PSCs. Results show mean ± SEM. *, $P < 0.05$, Mann-Whitney test. **(G)** Dot plot visualization of the scaled average expression of *Tnf* in malignant cells of KPC$^{Smad4\text{-WT}}$ (n=3) or KPC$^{Smad4\text{-KO}}$ (n=6) PDAC tumors, as analyzed by scRNA-seq. The color intensity represents the expression level and the size of the dots represents the percentage of expressing cells. **(H)** UMAP plots of malignant cells from KPC$^{Smad4\text{-WT}}$ or KPC$^{Smad4\text{-KO}}$ tumors colored by the normalized expression score of the TGF-β signaling HALLMARK pathway.

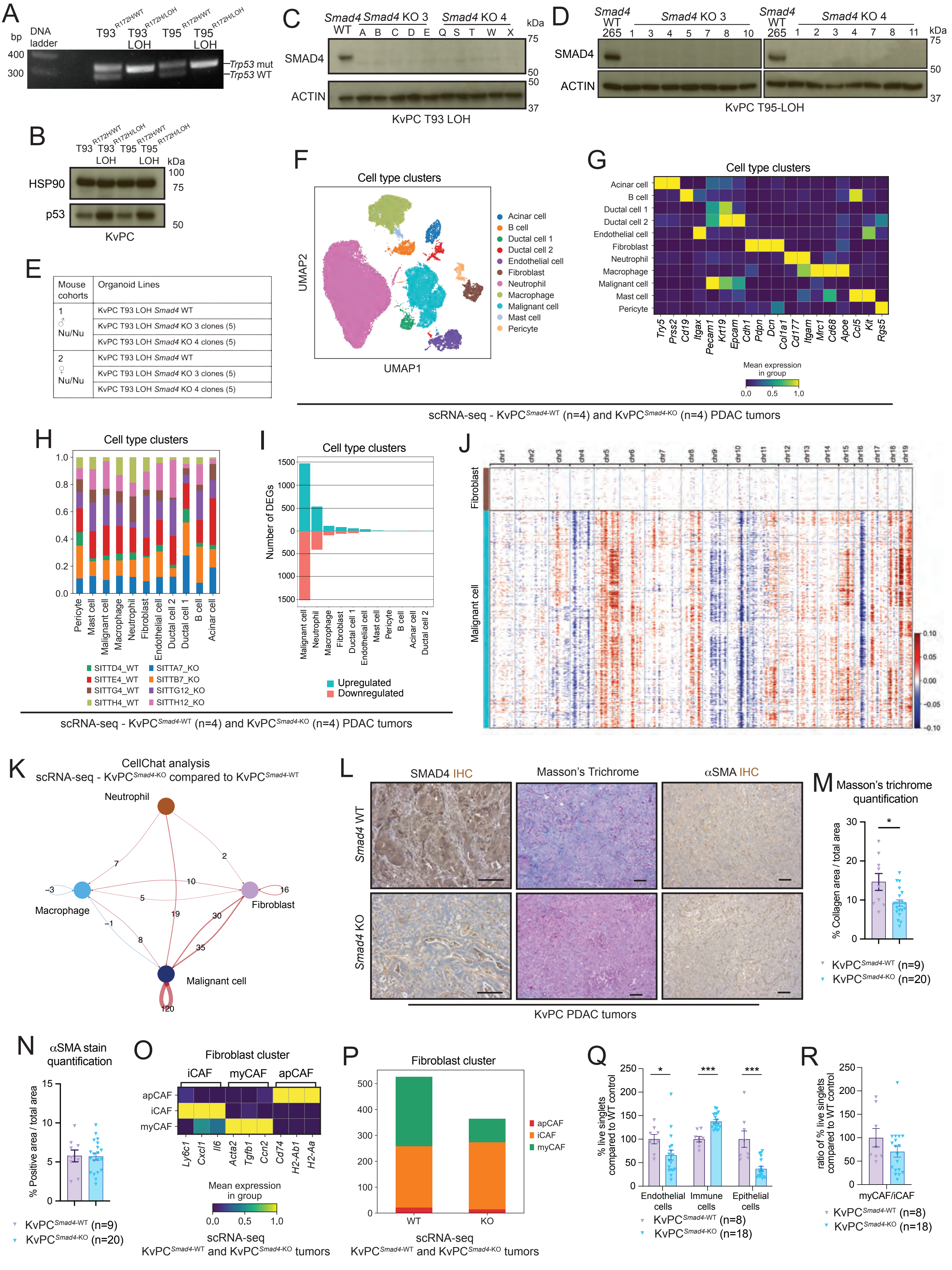
A
bp
DNA ladder
400
300
T93 R172H/WT
T93 LOH R172H/LOH
T95 R172H/WT
T95 LOH R172H/LOH
Trp53 mut
Trp53 WT
B
T93 R172H/WT
T93 LOH R172H/LOH
T95 R172H/WT
T95 LOH R172H/LOH
kDa
HSP90
100
75
p53
50
KvPC
C
Smad4 WT
Smad4 KO 3
A B C D E
Smad4 KO 4
Q S T W X
kDa
SMAD4
ACTIN
75
50
50
37
KvPC T93 LOH
D
Smad4 WT 265
Smad4 KO 3
1 3 4 5 7 8 10
Smad4 WT 265
Smad4 KO 4
1 2 3 4 7 8 11
kDa
SMAD4
ACTIN
75
50
37
KvPC T95-LOH
E
Mouse cohorts
Organoid Lines
1 ♂ Nu/Nu
KvPC T93 LOH Smad4 WT
KvPC T93 LOH Smad4 KO 3 clones (5)
KvPC T93 LOH Smad4 KO 4 clones (5)
2 ♀ Nu/Nu
KvPC T93 LOH Smad4 WT
KvPC T93 LOH Smad4 KO 3 clones (5)
KvPC T93 LOH Smad4 KO 4 clones (5)
F
Cell type clusters
UMAP2
UMAP1
Acinar cell
B cell
Ductal cell 1
Ductal cell 2
Endothelial cell
Fibroblast
Neutrophil
Macrophage
Malignant cell
Mast cell
Pericyte
G
Cell type clusters
Acinar cell
B cell
Ductal cell 1
Ductal cell 2
Endothelial cell
Fibroblast
Neutrophil
Macrophage
Malignant cell
Mast cell
Pericyte
Try5 Prss2 Cd19 Itgax Pecam1 Krt19 Epcam Cdh1 Pdpn Dcn Col1a1 Cd177 Itgam Mrc1 Cd68 Apoe Ccl5 Kit Rgs5
Mean expression in group
0.0 0.5 1.0
scRNA-seq - KvPC Smad4-WT (n=4) and KvPC Smad4-KO (n=4) PDAC tumors
H
Cell type clusters
1.0
0.8
0.6
0.4
0.2
0.0
Pericyte
Mast cell
Malignant cell
Macrophage
Neutrophil
Fibroblast
Endothelial cell
Ductal cell 2
Ductal cell 1
B cell
Acinar cell
SITTD4_WT
SITTE4_WT
SITTG4_WT
SITTH4_WT
SITTA7_KO
SITTB7_KO
SITTG12_KO
SITTH12_KO
I
Cell type clusters
1500
1000
500
0
500
1000
1500
Number of DEGs
Malignant cell
Neutrophil
Macrophage
Fibroblast
Ductal cell 1
Endothelial cell
Mast cell
Pericyte
B cell
Acinar cell
Ductal cell 2
Upregulated
Downregulated
scRNA-seq - KvPC Smad4-WT (n=4) and KvPC Smad4-KO (n=4) PDAC tumors
J
chr1 chr2 chr3 chr4 chr5 chr6 chr7 chr8 chr9 chr10 chr11 chr12 chr13 chr14 chr15 chr16 chr17 chr18 chr19
Fibroblast
Malignant cell
0.10
0.05
0.00
−0.05
−0.10
K
CellChat analysis
scRNA-seq - KvPC Smad4-KO compared to KvPC Smad4-WT
Neutrophil
Macrophage
Fibroblast
Malignant cell
7
2
10
5
−3
16
19
30
−1
8
35
120
L
SMAD4 IHC
Masson's Trichrome
αSMA IHC
Smad4 WT
Smad4 KO
KvPC PDAC tumors
M
Masson's trichrome quantification
% Collagen area / total area
30
20
10
0
*
KvPC Smad4-WT (n=9)
KvPC Smad4-KO (n=20)
N
αSMA stain quantification
% Positive area / total area
15
10
5
0
KvPC Smad4-WT (n=9)
KvPC Smad4-KO (n=20)
O
Fibroblast cluster
iCAF
myCAF
apCAF
apCAF
iCAF
myCAF
Ly6c1 Cxcl1 Il6 Acta2 Tgfb1 Ccn2 Cd74 H2-Ab1 H2-Aa
Mean expression in group
0.0 0.5 1.0
scRNA-seq
KvPC Smad4-WT and KvPC Smad4-KO tumors
P
Fibroblast cluster
500
400
300
200
100
0
WT
KO
apCAF
iCAF
myCAF
scRNA-seq
KvPC Smad4-WT and KvPC Smad4-KO tumors
Q
% live singlets compared to WT control
250
200
150
100
50
0
*
***
***
Endothelial cells
Immune cells
Epithelial cells
KvPC Smad4-WT (n=8)
KvPC Smad4-KO (n=18)
R
ratio of % live singlets compared to WT control
250
200
150
100
50
0
myCAF/iCAF
KvPC Smad4-WT (n=8)
KvPC Smad4-KO (n=18)

Figure S5

**Figure S5. *Smad4* loss drives a fibro-inflammatory stroma in KvPC PDAC**. **(A)** DNA gel showing *Trp53* status in T93 and T95 KvPC PDAC organoids with or without p53 loss of heterozygosity (LOH). WT, wild-type. mut, mutant. **(B)** Western blot analysis of p53 in T93 and T95 KvPC PDAC organoids with or without p53 LOH. HSP90, loading control. **(C-D)** Validation of *Smad4* KO KvPC PDAC organoids by western blot analysis of SMAD4 in KvPC$^{\mathit{Smad4}\text{-WT}}$ (i.e. *Rosa26* KO) or KvPC$^{\mathit{Smad4}\text{-KO}}$ clones cultured in complete organoid media. ACTIN, loading controls. **(E)** Table summarizing the experimental cohorts of orthotopic transplantation models of KvPC PDAC organoids in nu/nu mice. **(F)** UMAP plot of all cell types from KvPC$^{\mathit{Smad4}\text{-WT}}$ (n=4) or KvPC$^{\mathit{Smad4}\text{-KO}}$ (n=4) PDAC tumors analyzed by scRNA-seq. Different cell type clusters are color coded. **(G)** Heatmap of scaled expression of cell type-specific markers in each cell cluster of KvPC$^{\mathit{Smad4}\text{-WT}}$ (n=4) or KvPC$^{\mathit{Smad4}\text{-KO}}$ (n=4) PDAC tumors, as analyzed by scRNA-seq. Data are scaled such that the cluster with the lowest average expression = 0 and the highest = 1 for each gene. **(H)** Tumor sample contribution in KvPC$^{\mathit{Smad4}\text{-WT}}$ (n=4) or KvPC$^{\mathit{Smad4}\text{-KO}}$ (n=4) tumors, represented as bar plots showing proportions of the different tumor samples in each cell cluster. **(I)** Upregulated and downregulated differentially expressed genes (DEGs) in each cell type identified by pseudobulk analysis from scRNAseq of KvPC$^{\mathit{Smad4}\text{-WT}}$ or KvPC$^{\mathit{Smad4}\text{-KO}}$ tumors. False discovery rate (FDR) < 0.05. **(J)** Heatmap showing large-scale CNV profile of the fibroblast and malignant cell clusters identified by scRNA-seq. The color coding represents the CNV level based on a sliding window of 250 gene expression. Amplifications are shown in red and deletions are shown in blue. The fibroblast cluster was used as reference cell cluster. **(K)** Cell-cell communication analysis using CellChat showing the number of connections lost (in blue) or gained (in red) between malignant cells, fibroblasts, macrophages and neutrophils in KvPC$^{\mathit{Smad4}\text{-KO}}$ tumors compared to KvPC$^{\mathit{Smad4}\text{-WT}}$ tumors, as assessed by scRNA-seq. **(L)** Representative SMAD4, Masson's trichrome and $\alpha$SMA stains in KvPC$^{\mathit{Smad4}\text{-WT}}$ or KvPC$^{\mathit{Smad4}\text{-KO}}$ tumors. Scale bars, 50 $\mu$m. **(M)** Quantification of Masson's trichrome stain in KvPC$^{\mathit{Smad4}\text{-WT}}$ or KvPC$^{\mathit{Smad4}\text{-KO}}$ tumors. Results show mean $\pm$ SEM from 2 separate experiments, each with 1 WT group and 2 groups of KO pools from 2 different guides. *, $P < 0.05$, Mann-Whitney test. **(N)** Quantification of $\alpha$SMA stain in KvPC$^{\mathit{Smad4}\text{-WT}}$ or KvPC$^{\mathit{Smad4}\text{-KO}}$ tumors. Results show mean $\pm$ SEM from 2 separate experiments, each with 1 WT group and 2 groups of KO pools from 2 different guides. No statistical difference was found, as calculated by Mann-Whitney test. **(O)** Heatmap of scaled expression of CAF subtype-specific markers in each CAF cluster of KvPC$^{\mathit{Smad4}\text{-WT}}$ or KvPC$^{\mathit{Smad4}\text{-}}$

$^{\text{KO}}$ tumors, as analyzed by scRNA-seq. Data are scaled such that the cluster with the lowest average expression = 0 and the highest = 1 for each gene. **(P)** CAF sub-cluster abundance in fibroblasts of KvPC$^{\textit{Smad4}\text{-WT}}$ or KvPC$^{\textit{Smad4}\text{-KO}}$ tumors, as assessed by scRNA-seq. **(Q)** Flow cytometric analysis of endothelial cells (CD31$^{+}$CD45$^{-}$), epithelial cells (CD45$^{-}$CD31$^{-}$EpCAM$^{+}$) and immune cells (CD45$^{+}$) from live singlets in KvPC$^{\textit{Smad4}\text{-WT}}$ or KvPC$^{\textit{Smad4}\text{-KO}}$ tumors. Results show mean ± SEM from 2 separate experiments, each with 1 WT group and 2 groups of KO pools from 2 different guides. *, $P < 0.05$, ***, $P < 0.001$, Mann-Whitney test. **(R)** myCAF/iCAF ratio from live singlets in KvPC$^{\textit{Smad4}\text{-WT}}$ or KvPC$^{\textit{Smad4}\text{-KO}}$ tumors. Results show mean ± SEM from 2 separate experiments, each with 1 WT group and 2 groups of KO pools from 2 different guides. No statistical difference was found, as calculated by Mann-Whitney test.

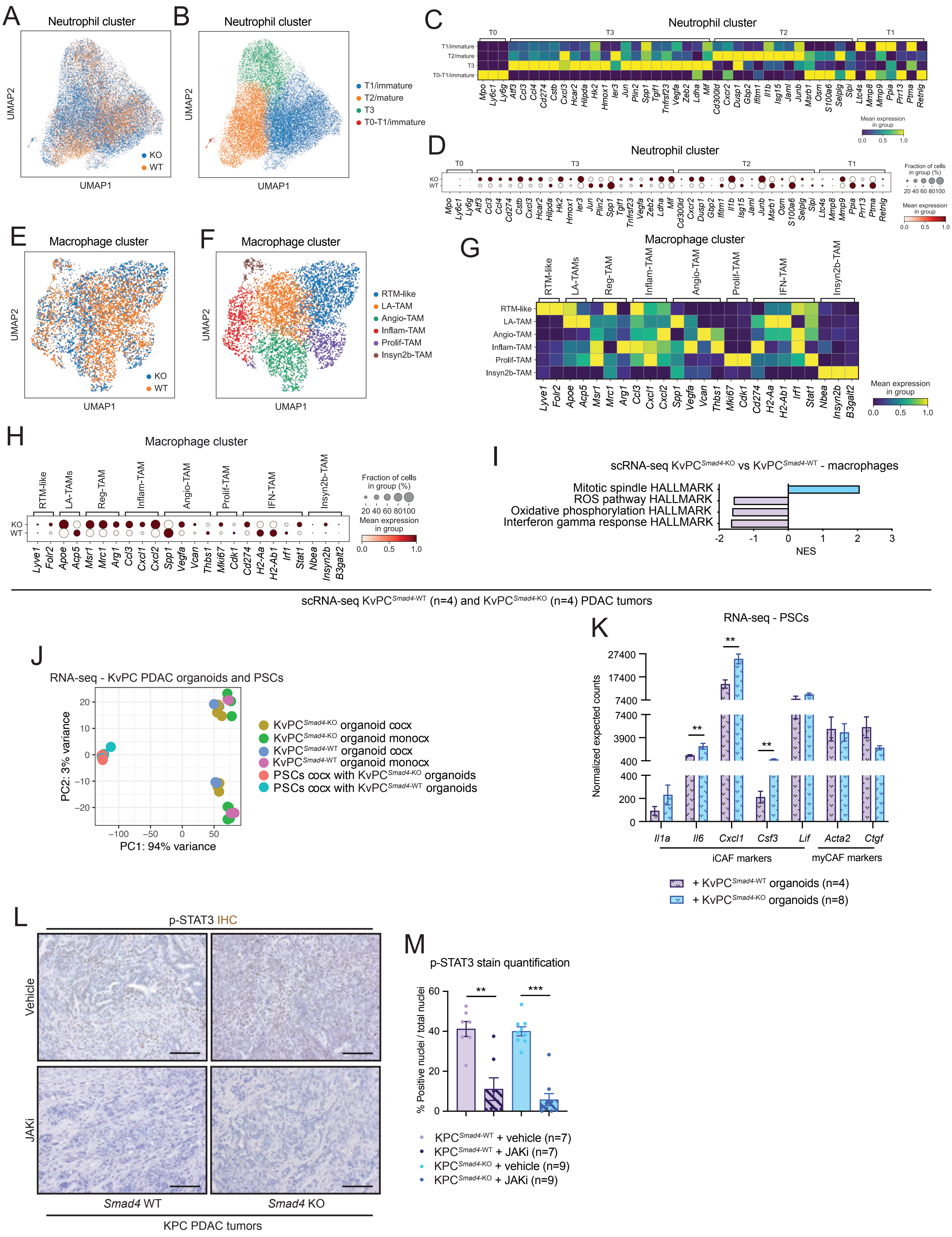
A
Neutrophil cluster
UMAP1
UMAP2
KO
WT
B
Neutrophil cluster
T1/immature
T2/mature
T3
T0-T1/immature
C
Neutrophil cluster
T0
T3
T2
T1
Mean expression in group
D
Neutrophil cluster
Fraction of cells in group (%)
Mean expression in group
Mpo Ly6c1 Ly6g Atf3 Ccl3 Ccl4 Cd274 Cstb Cxcl3 Hcar2 Hilpda Hk2 Hmox1 Ier3 Jun Plin2 Spp1 Tgif1 Tnfrsf23 Vegfa Zeb2 Ldha Mif Cd300ld Cxcr2 Dusp1 Gbp2 Ifitm1 Il1b Isg15 Jaml Junb Msrb1 Osm S100a6 Selplg Slpi Ltc4s Mmp8 Mmp9 Ppia Prr13 Ptma Retnlg
E
Macrophage cluster
KO
WT
F
Macrophage cluster
RTM-like
LA-TAM
Angio-TAM
Inflam-TAM
Prolif-TAM
Insyn2b-TAM
G
Macrophage cluster
RTM-like
LA-TAMs
Reg-TAM
Inflam-TAM
Angio-TAM
Prolif-TAM
IFN-TAM
Insyn2b-TAM
Mean expression in group
Lyve1 Folr2 Apoe Acp5 Msr1 Mrc1 Arg1 Ccl3 Cxcl1 Cxcl2 Spp1 Vegfa Vcan Thbs1 Mki67 Cdk1 Cd274 H2-Aa H2-Ab1 Irf1 Stat1 Nbea Insyn2b B3galt2
H
Macrophage cluster
Fraction of cells in group (%)
Mean expression in group
I
scRNA-seq KvPC Smad4-KO vs KvPC Smad4-WT - macrophages
Mitotic spindle HALLMARK
ROS pathway HALLMARK
Oxidative phosphorylation HALLMARK
Interferon gamma response HALLMARK
NES
scRNA-seq KvPC Smad4-WT (n=4) and KvPC Smad4-KO (n=4) PDAC tumors
J
RNA-seq - KvPC PDAC organoids and PSCs
PC1: 94% variance
PC2: 3% variance
KvPC Smad4-KO organoid cocx
KvPC Smad4-KO organoid monocx
KvPC Smad4-WT organoid cocx
KvPC Smad4-WT organoid monocx
PSCs cocx with KvPC Smad4-KO organoids
PSCs cocx with KvPC Smad4-WT organoids
K
RNA-seq - PSCs
Normalized expected counts
Il1a Il6 Cxcl1 Csf3 Lif Acta2 Ctgf
iCAF markers
myCAF markers
+ KvPC Smad4-WT organoids (n=4)
+ KvPC Smad4-KO organoids (n=8)
L
p-STAT3 IHC
Vehicle
JAKi
Smad4 WT
Smad4 KO
KPC PDAC tumors
M
p-STAT3 stain quantification
% Positive nuclei / total nuclei
KPC Smad4-WT + vehicle (n=7)
KPC Smad4-WT + JAKi (n=7)
KPC Smad4-KO + vehicle (n=9)
KPC Smad4-KO + JAKi (n=9)

Figure S6

**Figure S6.** ***Smad4*** **loss tunes signaling dependencies in PDAC with distinct KRAS status.** **(A-B)** UMAP plots of neutrophils from KvPC$^{Smad4\text{-WT}}$ (n=4) or KvPC$^{Smad4\text{-KO}}$ (n=4) PDAC tumors analyzed by scRNA-seq. Different genotypes **(A)** or neutrophil sub-clusters **(B)** are color coded. **(C)** Heatmap of scaled expression of neutrophil markers in distinct neutrophil sub-clusters from KvPC$^{Smad4\text{-WT}}$ and KvPC$^{Smad4\text{-KO}}$ tumors analyzed by scRNA-seq. Data are scaled such that the cluster with the lowest average expression = 0 and the highest = 1 for each gene. **(D)** Dot plot visualization of the scaled average expression of neutrophil markers in neutrophils from KvPC$^{Smad4\text{-WT}}$ or KvPC$^{Smad4\text{-KO}}$ tumors analyzed by scRNA-seq. The color intensity represents the expression level and the size of the dots represents the percentage of expressing cells. **(E-F)** UMAP plots of macrophages from KvPC$^{Smad4\text{-WT}}$ (n=4) or KvPC$^{Smad4\text{-KO}}$ (n=4) tumors analyzed by scRNA-seq. Different genotypes **(E)** and macrophage sub-clusters **(F)** are color coded. **(G)** Heatmap of scaled expression of macrophage markers in distinct macrophage sub-clusters from KvPC$^{Smad4\text{-WT}}$ and KvPC$^{Smad4\text{-KO}}$ tumors analyzed by scRNA-seq. Data are scaled such that the cluster with the lowest average expression = 0 and the highest = 1 for each gene. **(H)** Dot plot visualization of the scaled average expression of macrophage markers in macrophages from KvPC$^{Smad4\text{-WT}}$ or KvPC$^{Smad4\text{-KO}}$ tumors analyzed by scRNA-seq. The color intensity represents the expression level and the size of the dots represents the percentage of expressing cells. **(I)** Selected significantly upregulated (i.e. NES > 1.50 and FDR < 0.25) and downregulated (i.e. NES < -1.50 and FDR < 0.25) pathways identified by GSEA of macrophages from KvPC$^{Smad4\text{-KO}}$ compared to KvPC$^{Smad4\text{-WT}}$ tumors, as assessed by pseudobulk analysis from the scRNA-seq dataset. **(J)** PCA of PSCs and organoids flow-sorted from monocultures or co-cultures of KvPC PDAC organoids with PSCs. **(K)** RNA-seq expression of iCAF and myCAF markers in PSCs flow-sorted from co-cultures with KvPC$^{Smad4\text{-WT}}$ or KvPC$^{Smad4\text{-KO}}$ PDAC organoids. Results show mean ± SEM. **, $P < 0.01$, Mann-Whitney test. **(L)** Representative p-STAT3 IHC stains in 2-week vehicle- or JAK inhibitor (JAKi, AZD1480)- treated KPC$^{Smad4\text{-WT}}$ and KPC$^{Smad4\text{-KO}}$ PDAC tumors. Scale bars, 50 μm. **(M)** Quantification of p-STAT3 IHC stain in 2-week vehicle- or JAKi-treated KPC$^{Smad4\text{-WT}}$ and KPC$^{Smad4\text{-KO}}$ PDAC tumors taken on the same day of the last dose. Results show mean ± SEM. **, $P < 0.01$; ***, $P < 0.001$, Mann-Whitney test.

#### SUPPLEMENTARY TABLES

**Table S1. Clinical information and characteristics of *SMAD4* WT and *SMAD4*-deficient human PDAC tissues, related to Figure 1.**

**Table S2. Clinical information and characteristics of *SMAD4* WT and *SMAD4*-deficient human PDAC organoids used for the generation of orthotopic transplantation models, related to Figure 1.**

**Table S3. Single-cell RNA-sequencing of murine KPC PDAC tumors – differential expression analysis and GSEA, related to Figures 2, 3 and 4.**

**Table S4. RNA-sequencing of PSCs flow-sorted from KPC PDAC organoid co-cultures – differential expression analysis, normalized expected counts and GSEA, related to Figure 3.**

**Table S5. RNA-sequencing of *Smad4* KO and *Smad4* WT KPC PDAC organoids flow-sorted from co-cultures with PSCs – differential expression analysis, normalized expected counts and GSEA, related to Figure 4.**

**Table S6. Ligand activity plots from NicheNet analyses showing the top ligands that regulate a group of targeted genes, related to Figure 4.**

**Table S7. Single-cell RNA-sequencing of murine KvPC PDAC tumors – GSEA, related to Figures 5 and 6.**

**Table S8. RNA-sequencing of PSCs flow-sorted from KvPC PDAC organoid co-cultures – differential expression analysis, normalized expected counts and GSEA, related to Figure 6.**

**Table S9. RNA-sequencing of *Smad4* KO and *Smad4* WT KvPC PDAC organoids flow-sorted from co-cultures with PSCs – differential expression analysis, normalized expected counts and GSEA, related to Figure 6.**
